# Imaging the structure and dynamic activity of retinal microglia and macrophage-like cells in the living human eye

**DOI:** 10.1101/2022.04.30.490173

**Authors:** Yuhua Rui, Daniel M.W. Lee, Min Zhang, Valerie C Snyder, Rashmi Raghuraman, Elena Gofas-Salas, Pedro Mecê, Sanya Yadav, Pavan Tiruveedhula, Kate Grieve, José-Alain Sahel, Marie-Hélène Errera, Ethan A. Rossi

## Abstract

**Purpose:** We recently showed how a refined sequential detection pattern and image processing pipeline for multi-offset adaptive optics scanning light ophthalmoscopy (AOSLO) can increase the contrast of weakly scattering inner retinal structures, including microglia. However, sequential detection was still time-consuming, preventing dynamics from being monitored over short intervals (< 3 mins). Here we show that simultaneous fiber-bundle (FB) detection can overcomes this limitation to reveal the structure and dynamic activity of microglia and macrophage-like cells in healthy and diseased retinae.

**Methods:** We designed and implemented a custom 7-fiber optical FB with one central confocal fiber and six larger fibers for multi-offset detection in AOSLO at a single focal plane. We imaged the ganglion cell layer at several locations at multiple timepoints (from minutes to weeks) in 8 healthy participants and in 4 patients with ocular infections or inflammation, including ocular syphilis and posterior uveitis. Microglia and immune cells were manually segmented to quantify cell morphometry and motility.

**Results:** Fiber-bundle detection reduced single acquisition time to 20-30 seconds, enabling imaging over larger areas and monitoring of dynamics over shorter intervals. Presumed microglia in healthy retinas had an average diameter of 12.8 μm and with a spectrum of morphologies including circular cells and elongated cells with visible processes. We also detected the somas of putative macroglia, potentially astrocytes, near the optic nerve head. Microglia moved slowly in normal eyes (0.02μm/sec, on average) but speed increased in patients with active infections or inflammation (up to 2.37μm/sec). Microglia activity was absent in a patient with chronic uveitis that was quiescent but apparent over short intervals in an active uveitis retina. In a patient with ocular syphilis imaged at multiple timepoints during treatment, macrophage-like cells containing granular internal structures were seen. Decreases in the quantity and motility of these immune cells were correlated with improvements to vision and other structural and systemic biomarkers.

**Conclusions:** FB-AOSLO enable simplified optical setup with easy alignment and implementation. We imaged the fine-scale structure and dynamics of microglia and macrophage-like cells during active infection and inflammation in the living eye for the first time. In healthy eyes we also detected putative glia cell near the optic nerve head. FB-AOSLO offers promise as a powerful tool for detecting and monitoring retinal inflammation and infection in the living eye over short response to treatment.

## 1. Introduction

Retina microglia constitute a prominent part of the resident glial population and are the dominant resident myeloid population^1^ along with perivascular macrophages. Microglia actively participate during development in the pruning and sculpting of the retinal cells and their circuitry^2^ and have been shown to be necessary for the healthy function of the mature retina, including through the maintenance of synapses in the plexiform layers^3^. In the healthy adult retina microglia exist in a regularly ordered ramified state^2^. In primates, their highest densities are observed in the inner and outer plexiform layers while a smaller proportion reside in the ganglion cell layer^4^. Ramified microglia have cell morphologies that are dendritic in appearance with many processes extending from the soma that constantly survey the local microenvironment.^5^ Resting microglia form a network of potential immuno-effector cells^6^ that can become activated during inflammation. Depending on the inflammatory stimuli, these cells can exhibit a wide range of actions and roles such as driving acute inflammatory responses and both initiating and regulating lymphocyte-driven adaptive responses. Resident myeloid cells (monocytes/macrophages) can trigger inflammatory responses to pathogens and tissue trauma and pave the way for the progressive recruitment of circulating myeloid cells such as neutrophils and inflammatory monocytes/macrophages and non-myeloid natural killer cells into the eye^7^.

Though previously inaccessible to direct cellular-level ophthalmic *in vivo* imaging, recent advances in adaptive optics ophthalmoscopy (AOO) have begun to reveal these cells and their dynamic activity with high resolution^5,8,9^. AOO permits high-resolution imaging of the human retina *in vivo* at a cellular level by compensating for the optical aberrations of the eye^10^. The addition of adaptive optics to multiple ophthalmic imaging modalities, such as adaptive optics scanning light ophthalmoscopy (AOSLO)^11^, adaptive optics optical coherence tomography (AOOCT)^12^, and flood illumination adaptive optics (FIAO)^13^ have enabled visualization of many of the cell classes within the living human eye, with some retinal cell classes (e.g. cones and retinal pigmented epithelial cells) accessible to visualization in multiple modalities. Over the past several years, the list of retinal cell classes accessible to *in vivo* imaging has continued to grow. In AOOCT, advances towards imaging of additional cell classes, such as retinal ganglion cells and macrophages, have come from higher speed devices with improved image processing and registration that enable the averaging of many volumetric datasets^5,14,15^. In AOSLO, additional cell classes and structures have recently become accessible mostly due to changes in how the light is detected.

Various forms of non-confocal AOSLO have provided access to weakly scattering retinal structures that previously were challenging to image using traditional confocal imaging. Unlike confocal imaging where a pinhole aperture is set on the optical axis at the retinal conjugate focal plane to reject light from out of focus layers of the retina, the principle behind non-confocal imaging modes is to purposely acquire the non-confocal portion of the light distribution that falls outside the confocal aperture detection area. It is thought that these techniques preferentially collect light that has been multiply scattered^16,17^. The first and simplest approach is to use an aperture that is purposely displaced from the optical axis^16,17^. Offset aperture detection in AOSLO was shown to enhance the contrast of certain retinal structures, such as capillary walls^16^. Since then, additional non-confocal AOSLO setups have been devised that differ mainly in the detection pattern and number of detectors used. Scoles et al. showed that splitting the non-confocal light distribution in two and directing it to different detectors yields two images of the same structures that could be combined to further enhance the contrast of weakly scattering structures, such as the inner segments of cones^18^. Rossi et al. showed that the non-confocal light distribution could be further sub-divided and similarly combined by using an offset aperture sequentially positioned at multiple points across the retinal conjugate focal to improve the contrast of weakly scattering cells, such as retinal ganglion cells^8,19^ and microglia^8^. Recently, it was shown that adding an additional orthogonal split to the split-detection setup to achieve 4 quadrant detection^20^ combined with an emboss filtering approach, could reveal a specific class of retinal microglia, the vitreous cortex hyalocytes, and the dynamics of their processes with high contrast.^9^ An optical model has been proposed that suggests that these methods enhance contrast by exploiting spatial variations in the refractive index in a similar way to phase contrast microscopy^21^.

We recently showed that multi-offset detection could be optimized for imaging human retinal ganglion cell layer structures through the use of an improved radial detection pattern^8^ and spatial-frequency based image fusion^22^. However, sequential multi-offset^8^ detection remains slow due to the need to physically move the aperture across the retinal conjugate focal plane and acquire an image sequence at each location. Total acquisition of one retinal location with 8 offsets and 20 second recordings was 3 minutes. This relatively long duration prevents large areas of the retina to be acquired in a single imaging session and dynamics from being detected over shorter intervals. Further, during each 3-minute acquisition, the AO correction and thus optical quality fluctuates, and eye movements shift the field of view, reducing the common area of overlap for subsequent image fusion.

We imaged both normal healthy retinae and inflammatory retinae of patients with uveitis, vasculitis, and ocular syphilis. Current commonly used clinical imaging modalities for managing posterior uveitis and particularly vasculitis include fluorescein angiography (FA), fundus autofluorescence imaging (FAF), and Optical Coherence Tomography (OCT)^23,24^. Only a few AO imaging case-reports and small case series have studied the photoreceptor layer and vasculature changes in uveitis but microglia have not been studied^25–27^. In this current study, we explore the ganglion cell layer at several locations at multiple timepoints (from minutes to weeks) in eyes affected by posterior uveitis to evaluate the efficacy of fiber bundle detection for imaging microglia and their motility in their activated state.

Here we show how the multi-offset detection scheme can be substantially improved through acquisition of all offsets simultaneously using an optical fiber bundle, after Mozaffari et al.^28^. This enabled improved detection of retinal ganglion cell layer structures including visualization of microglia and macrophage-like cells and their movement across short timescales. In this current study, we imaged normal healthy eyes and compared them to patients with uveitis, vasculitis, and ocular syphilis, and we evaluated the efficacy of fiber bundle detection for imaging microglia and macrophage-like cells and their motility in their activated state.

## 2. Materials and Methods

### 2.1 AOSLO system and Fiber Bundle based detection pathway

The Pittsburgh AOSLO, described in detail elsewhere^8,29^, was modified for these experiments; only relevant system details are reported here. The light detection pathway and data acquisition system was modified for fiber bundle detection but the rest of the system was configured as previously described^8,29^. Two light sources were used, a 909 nm laser diode (QFLD-905-200S, Qphotonics, Ann Arbor MI, USA) for wavefront sensing, and a 795 nm (FWHM = 15 nm) super-luminescent diode that scanned in a raster fashion at a frame rate of 30 Hz for illuminating a 1.5°× 1.5° field of view (FOV) of the retina. Peak power at the cornea for the light sources was 590 μW (795 nm), and 21 μW (909 nm); these light levels are below the limits specified by the ANSI^30^. Figure.1A shows the AOSLO system with a blue box highlighting the fiber bundle detection pathway. Light backscattered from the retina was directed to the fiber bundle using a dichroic mirror that reflects everything below 810 nm (T810lpxr, Chroma, USA) and was then focused onto the fiber bundle with an achromatic doublet (AC254-100-B, Thorlabs GmbH, Germany).

**Figure 1.**
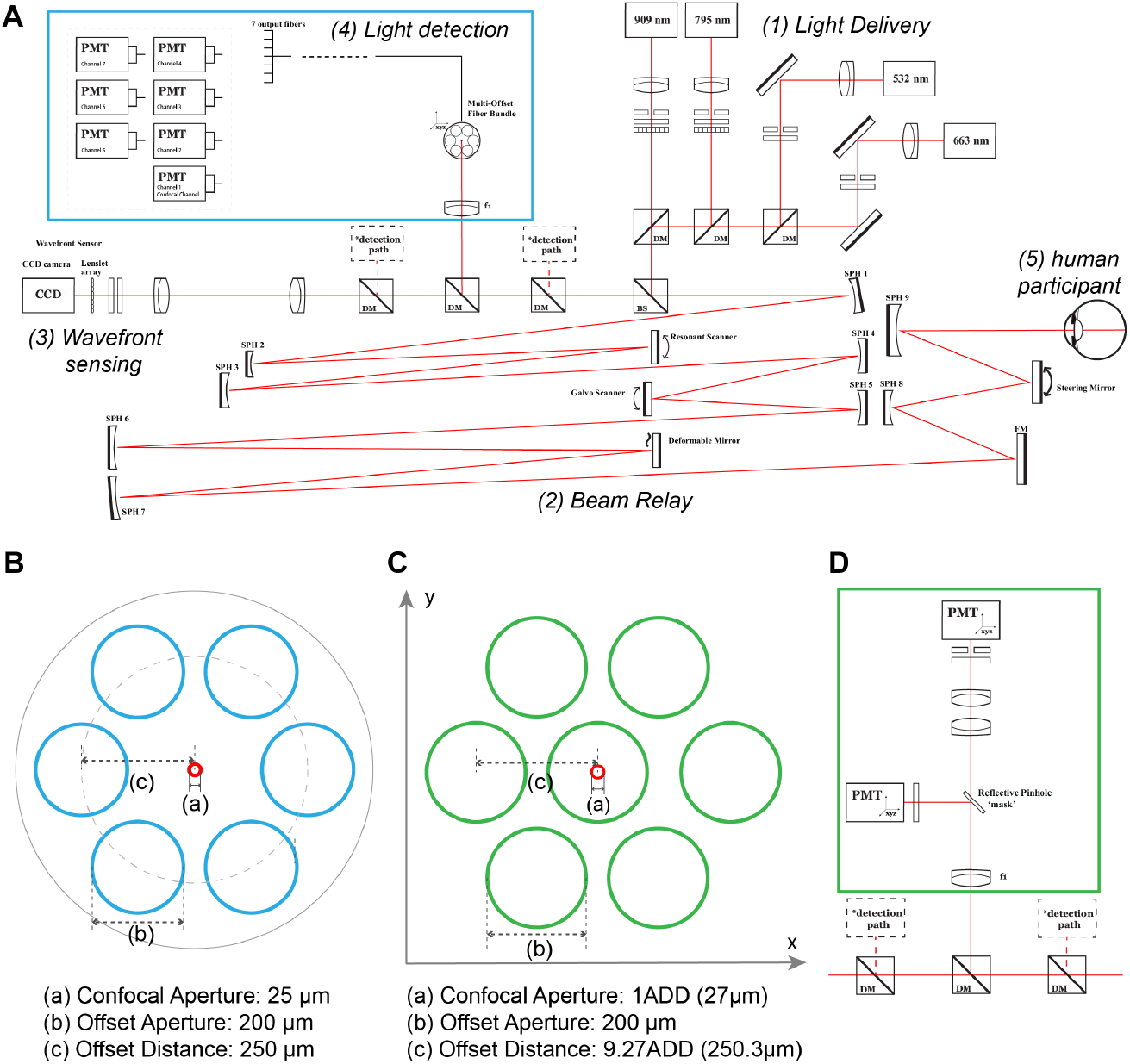
Schematic Drawing of Pittsburgh AOSLO system and Fiber Bundle detection design. **(A)** optical layout of the Pittsburgh AOSLO, modified from initial system setup described previously^8^. **(B)** geometric arrangement of fibers on Fiber Bundle detection side, distance (a) represents confocal aperture diameter, (b) represents offset aperture, and (c) represents offset distance. Dashed circle is only for representation purpose, it connects the center of offset apertures, indicating the radial layout of offset apertures. **(C)** radial pattern set in sequential multi-offset imaging with parameters as close to FB design as possible for comparing image quality from two different imaging approaches. Distance (a-c) represent the same measurements as in (B). **(D)** detection section of sequential multi-offset imaging^8^.

We designed the fiber bundle (FB) in-house and had it custom built to our specifications by FiberTech Optica Inc., (Kitchener, ON, Canada). The FB was mounted on a motorized translation stage to allow for precise computer controlled alignment^8^. Fig.1B shows the geometric arrangement of the fibers on the detection side. The six radially arranged apertures (blue circles in Fig.1B) mimics the pattern that we found to be most effective in our previous sequential multi-offset study^8^; Though in our sequential work we used 8 apertures, we reduced this to 6 for the FB and made some other slight modifications to conform to manufacturing constraints based on the optical fibers that were available for FB fabrication. The FB implementation is much simpler than our previous sequential multi-offset detection setup that is shown in Fig.1D because it eliminates the need for the reflective pinhole mask that was needed to direct the confocal light to a first PMT and removes the additional lens telescope that was needed to generate a second retinal conjugate focal plane^8^.

The Airy Disk Diameter (ADD) at the retinal conjugate focal plane was 27 μm. The central aperture of the FB served as the pinhole for confocal imaging and has a diameter of 25 μm (0.93 ADD). The six larger surrounding apertures for offset detection have a diameter of 200 μm (7.4 ADD) and are centered at 250 μm (9.3 ADD) from the center of the confocal fiber. Light detected by the FB fanned out into 7 individual fibers that were each directly connected to their own PMT detector (H7422-50, Hamamatsu, Inc.) using SMA to C-mount adapters with no additional optical elements. Data acquisition from the FB required the ability to digitize 7 PMT signals simultaneously. To accomplish this, we utilized two legacy frame grabber boards (Helios eA/XA; HEL 2M QHAL E*; Matrox, Montreal, QC, Canada). Each Helios board is capable of digitizing four analog signals simultaneously and we synchronized two boards using custom electronics and software to acquire up to a total of 8 simultaneous channels, only 7 of which were utilized in this study.

### 2.2 Participants

This study was approved by the Institutional Review Board of the University of Pittsburgh and adhered to the tenets of the Declaration of Helsinki. Twelve participants were enrolled in the study and their demographic information is shown in Table 1. Written informed consent was acquired prior to enrollment for each participant following a detailed explanation of experimental procedures both verbally and in writing. Before imaging, one drop of 1% tropicamide and one drop of 2.5% phenylephrine hydrochloride were applied for pupil dilation.

**Table 1.** Participant demographics.

| Category | Participants<br>(n) | Sex | | Eye imaged | | Age<br>(mean ± SD) | Axial Length<br>(mean ± SD) |
| --- | --- | --- | --- | --- | --- | --- | --- |
|  |  | Male | Female | OD | OS |  |  |
| 1. healthy retina (P) | 8 | 5 | 3 | 7 | 1 | 30 ± 6 | 24.36 ± 0.54 |
| 2. retinal<br>inflammation (Pt) | 4 | 1 | 3 | 3 | 1 | 51 ± 13 | 23.88 ± 0.83 |

The study consisted of two parts, in part one we imaged healthy eyes to evaluate FB detection, compare it to sequential multi-offset imaging, and evaluate microglia imaging in eyes with healthy retinae; in part two we imaged patients with posterior segment inflammation. Eight healthy participants (n= 8 eyes) imaged were all free from ocular conditions including inherited eye diseases, retinal condition, uveitis, or glaucoma (hereafter referred to as P1-P8). Four patients recruited from the UPMC ophthalmology clinics (hereafter referred as Pt1-Pt4) were affected by uveitis, including cases of acute uveitis infectious including acute syphilitic posterior placoid chorioretinitis (ASPPC) (n=1 eye, 1 patient) and active idiopathic posterior uveitis (n=1 eye, 1 patient), chronic active (n=1 eye, 1 patient) and quiescent chronic idiopathic posterior uveitis (n=1 eye, 1 patient). Medical history and treatment information for the four patients are shown in Table 2.

**Table 2.** Case Description of uveitis patients.

| patient No. | sex | age | eye | major diagnosis | imaging sessions | duration of uveitis at initial imaging | duration of treatment at imaging | cClinical features | fluorescein angiography (FA) |
| --- | --- | --- | --- | --- | --- | --- | --- | --- | --- |
| 1 | Male | 52 | OS | ASPPC | 3 | 6 days | #1: 6 days (penicillin IM)<br>#2: 26 days<br>#3: 41 days | granular dots within the foveal area | N/A |
| 2 | Female | 69 | OD | acute PU | 1 | 3 months | 2 weeks (oral steroids) | N/A | Improvement of retinal vasculitis as compared to May 2021 OD. |
| 3 | Female | 35 | OD | chronic PU, CSME | 2 | 8 years | #1: 5 months (ADA), 3 months (PSTA)<br>#2: 12 months (ADA), 6 months (trisenec) | Peripheral ischemia, no vessel sheathing | #1: mid periphery vessels leakage, diffuse capillary staining<br>#2: mid periphery vessels leakage, diffuse capillary staining, unchanged |
| 4 | Female | 48 | OD | chronic PU, CSME | 1 | 20 months | 7 months (PSTA) | Peripheral ischemia, no vessel sheathing | widespread capillary staining, peripapillary nasal vessels leakage |

ASPPC: acute placoid chorioretinitis; PU: posterior uveitis; CSME: cystoid macular edema; IM: intramuscular; ADA: adalimumab; PSTA: Posterior Subtenon injection of Triamcinolone Acetonide; N/A: not available.

### 2.3 Imaging protocols

#### Clinical imaging

An infrared reflectance confocal SLO (cSLO) image (Spectralis HRA, Heidelberg Engineering, Inc) was acquired on all participants for orientation and navigation during AOSLO imaging. All participants also had their axial length measured (IOLmaster; Carl Zeiss Meditec) to calculate the retinal magnification and scale the images across modalities; images were scaled using the Gullstrand simplified relaxed eye model, as previously described^8^. Select participants were imaged using a commercial flood-illumination adaptive optics (FIAO) device (device version: rtx1-e; software version: AOImage 3.4; Imagine Eyes, Orsay, France). Patients underwent additional clinical imaging as part of their standard of care, including spectral domain optical coherence tomography (SD-OCT; Spectralis HRA+OCT; Heidelberg Engineering, Heidelberg, Germany) and fluorescein angiography. (Spectralis HRA+OCT; Optos Wide Field (UWF) Optomap, Optos, USA).

#### Comparing FB-AOSLO to sequential multi-offset

To compare the retinal images obtained from the FB to our previous sequential setup, we carried out a series of imaging sessions with the sequential multi-offset setup before installing the FB using the detection pattern shown in Figure 1(C). As noted above, there were some slight differences between the sequential and FB pattern, but we consider these to be negligible for the purposes of this comparison. After installing the fiber bundle, we also compared FB images acquired with different exposure times (from 20–60s). Three participants were imaged at 6 different retinal locations; anticipating some potential challenges in imaging the exact same focal planes across the different imaging configurations, we imaged 2 different retinal layers at each retinal location for both sequential and FB imaging. Image sequences were acquired with an exposure time of 30 seconds per aperture position, resulting in nearly 4 minutes for one sequential acquisition but only 30 seconds for each corresponding FB recording. With the extra time available for FB imaging, we also evaluated the capability to acquire multiple images at additional locations to form a larger montage. For FB imaging, we used the steering system of the AOSLO to rapidly image across multiple retinal locations; we used a separation of 1 degree between adjacent locations to give enough overlap for later montaging. Two regions were imaged: temporal to the fovea (6 participants), and around the optic nerve head region (3 participants). During image acquisition, we used the deformable mirror (DM) to focus on different retinal planes. Using the DM we first set the focus to a plane where the nerve fiber bundles were high contrast in the confocal channel and then displaced the focus slightly below this plane where the capillaries start to show high contrast in offset channels in an attempt to focus near the retinal ganglion cell layer. We kept imaging at this layer throughout the study for both healthy participants and uveitis patients. During imaging across multiple locations (e.g., for montage creation) only minor adjustments in focus were made at the border of the montage region for better image quality. Near the optic nerve head, we evaluated FB detection at a few additional focal planes from the RNFL to the photoreceptor layer.

#### Imaging of retinal microglia and their dynamics in healthy eyes

We imaged multiple retinal locations to form a larger montage for characterizing cell morphology in 6 participants; an average of 16 locations were stitched together in Adobe Photoshop (Adobe Systems Inc., San Jose, CA, USA) to generate a larger montage. To potentially detect movement of microglia, we imaged several locations repeatedly during each imaging session. The focal plane was fixed and the interval between acquisitions varied to observe motility over a variety of intervals from seconds up to 30 minutes.

#### Monitoring microglia in patients with inflammation

We evaluated FB detection in 4 patients with retinal infection and inflammation at varying intervals. Imaging of the ASPPC participant (Pt1) was directed to a position near one of the largest plaques seen on SLO near the fovea (Fig.6A). Cells were readily visible even in the live FB images and we used this to guide focus to the plane where we could detect the greatest number of cells. We imaged this location multiple times on the first imaging session to assess cell motility by forming time-lapse image sequences. We re-imaged his retina at the same location and approximate focal plane on follow-up 20 and 35 days later to evaluate changes in response to treatment. At each timepoint we acquired multiple images on the same location to assess cell motility. For three other patients with posterior uveitis, we selected the retinal location for imaging based on the clinical images (including FA) and clinical evaluation on the disease to visualize microglial changes with uveitis. We focused on the ganglion cell layer and at certain locations performed multiple acquisition to evaluate cell motility.

### 2.4 Image acquisition and processing

After pupil dilation patients were seated in front of the AOSLO and we used a custom-designed forehead and chin rest mounted on a motorized stage for pupil alignment. The AOSLO steering mirror sub-system and integrated fixation targeting system allowed for precise positioning of the AOSLO imaging field across the retina. We gave participants several breaks between acquisitions and kept the total imaging session duration (include pupil dilation and breaks) to no longer than 90 minutes.

Image sequences were processed using several custom MATLAB scripts (R2020a, The MathWorks, Inc., Natick, MA). Sinusoidal artifacts in all videos were corrected using previously published methods^31^. Confocal image sequences were registered using our custom-developed strip-based registration algorithm^32^ and the motion correction was applied to each offset channel for co-registration. Average images for each offset position were generated (hereafter called ‘offset images’). Offset images went through linear combination and spatial frequency based image reconstruction (SMART) processing, as described in detail previously^22^ to enhance image contrast; hereafter these will be referred to as ‘FB images’ (i.e. Fiber Bundle multi-offset images).

For the majority of the imaging study, FB images were generated from 30-second image sequences whereas for the ASPPC patient (Pt1), considering the acute phase of this ocular infection and the rapid cell motility observed, we kept the image acquisition 30 seconds but when processing afterwards, each acquisition was break into 3 sets of 10-seconds image sequences and therefore generated corresponding FB images.

### 2.5 Image analysis

When comparing image quality between the FB and sequential imaging approaches as well as assessing images obtained at different exposure durations with FB imaging, we evaluated the power distribution within spatial frequencies corresponding to cell mosaics of interest. Radially averaged power spectral density (PSD) was generated in MATLAB and the power was normalized and plotted in a log scale to display the normalized PSD across spatial frequencies (in cycles per mm).

To characterize the microglia, we generated several images, including: 1) perfusion maps for comparing with vasculature, 2) retinal montages for cell morphometric quantification, and 3) time-lapse image sequences for quantification of cell motility. For each acquisition, perfusion maps were first generated from each of the 6 individual offset image sequences and then averaged together to form a final perfusion map with higher contrast. Each Perfusion map was generated from registered image sequence of 900 frames. We first applied a spatial filter to remove the low spatial frequency components (cutoff at 0.02) and then a median filter with 1-by-1 neighborhood to smooth the potential noise on every frame and then computed the variance across all frames throughout the sequence to reveal the motion contrast caused by blood flow in perfused vessels. Retinal montages of the FB images and perfusion maps were generated using Photoshop. Level, contrast, and brightness of the individual images were manually adjusted to have consistency across the different imaging locations. Hard borders between adjacent images were minimized to optimize visualization and quantification by feathering the boundaries of adjacent montage images using the eraser tool with a gaussian shaped brush in Photoshop. For visualizing microglial cell motility, FB images from same retinal location but different acquisition times were registered using sub-pixel precision^33^ and cropped to a mutual region. Time-lapse videos were then generated in ImageJ^34^.

Microglia were manually segmented to quantify morphometry. Two types of images were analyzed: 1) the 6 large retinal montages were analyzed to characterize cell morphology, and 2) every single frame in each time-lapse sequence were analyzed to evaluate motility. Independent labeling of microglia and macrophage-like cells was done for both montages (3 graders) and time-lapse videos (2 graders). Graders were asked to trace the cell boundaries for all the cells they can see using the pencil tool with a one-pixel sized brush in Adobe Photoshop. Labeled images from different graders were manually compared and cells that were only marked by one grader went through an additional round of determination by all three graders where a consensus was used to decide whether to include or reject the cells only marked by one of the three graders. This step ensured that we included all the cells detected by at least two graders. Labeled images were then converted to a binary image, and measurements were extracted with MATLAB including their size (represented by the cell area), major and minor axis (represented by computing the best-fitting ellipse for the label using MATLAB function *regionprops),* and positions (X, Y coordinates). Due to the elliptical shape of most cells, we also computed eccentricity (i.e., the minor axis divided by the major axis) to quantify their morphology as more circular or more elliptical (Fig.5B). Eccentricity ranges from 0–1, with more rounded cells closer to 1 (i.e., an eccentricity equal to 1 is a perfect circle) and more elongated cells closer to 0. The average of the major and minor axis was taken to represent the cell diameter. The cartesian coordinates were used to measure the cell displacement between different observation time points and divided by the time interval to get their speed (in μm/sec).

## 3. Results

### 3.1 Fiber Bundle detection generates high quality retinal images with increased efficiency and additional capabilities

We compared images of the same retinal location obtained from FB detection to those from sequential multi-offset Fig.2(A-D) with nearly identical detection patterns (Fig.1 B and C). Both methods generated retinal images with high contrast and no visible difference in image quality. Normalized radially averaged PSDs are plotted in Fig.2(I), the highlighted region denotes the spatial frequency band that is presumed for the retinal ganglion cell mosaic at this retinal eccentricity (80-100 cycle/mm). Both FB images have higher energy than sequential images in the spatial frequency range of interest. At high spatial frequencies (>300 cycle/mm), FB images showed lower energy than sequential images likely due to reduced noise. An additional example comparing FB and sequential imaging is shown in SFig.1. Fig.2(E-H) show FB images of the same retinal location but with different exposure times; the corresponding exposure times are marked on the image (note Fig.2(A) shows a 30s exposure). Fig.2(J) shows the radially averaged PSD plot for images Fig.2(A, E-H). In the spatial frequency range of interest for detecting RGCs, a similar energy level was observed, with 20s duration images showing only a modestly higher energy in this band than the rest. Based on these results, a 30 second exposure time was used for all subsequent FB imaging.

**Figure 2.**
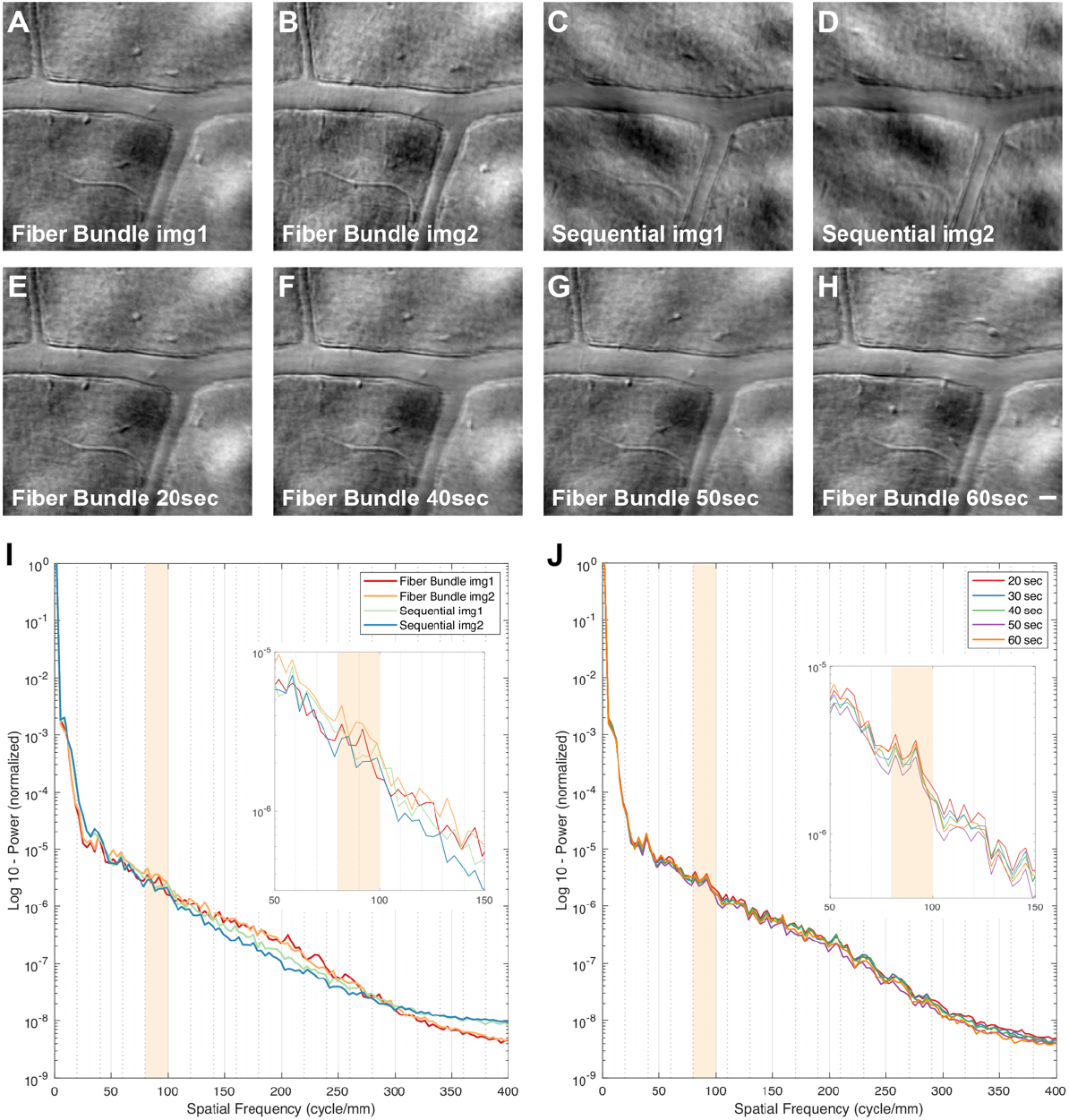
Comparison between FB and sequential multi-offset imaging and evaluation of exposure duration on FB images. **(A-D)** images of 2 different focal planes for both fiber bundle (FB) and sequential detection acquired with 30s recordings. **(E-H)** FB images of retinal location at the same focal plane as (A) with different exposure time. Scale bar is 20 μm and applied to image (A-H). **(I)** normalized radially averaged PSD log plot of images (A-D). Orange box highlights the spatial frequency band expected for retinal ganglion cells at this eccentricity. Inset plot shows zoomed in view of spatial frequency range from 50–150 cycle/mm. **(J)** normalized radially averaged PSD log plot of images (A, E-H). Orange box highlights spatial frequencies of interest. Inset plot shows zoomed in view of spatial frequency range from 50–150 cycle/mm.

The reduced acquisition time permitted multiple retinal locations to be imaged in a single session. Multiple FB montages were generated to reveal retinal structure across larger regions (Fig.4A and SFig.3A). FB detection also enabled imaging of some retinal areas that previously had been challenging to image with sequential detection including the optic nerve head (ONH) region. Fixational eye movements can make the ONH and surrounding retina challenging to image as the eye moves the field of view across substantially different axial planes; this can cause the AO correction to sometimes be unstable, especially over the timescales of a few minutes that had been required for sequential detection. We successfully imaged around the optic nerve head region using FB detection in 3 healthy participants (Fig.3 and SFig.2). The focal plane for image acquisition was achieved by setting the defocus on the deformable mirror, and we used the relative value and image appearance to localize the retinal depth. Several cell-like structures were seen along the optic rim in FB images that were invisible in confocal channel (Fig.3B). These cell-like structures appear to have a relatively uniform circular appearance across their internal surface. A total of 15 of these cells were detected and measured on the optic rim from two participants (Fig.3 and SFig.2), their average diameter was 23 μm (range 18-28μm) and they were seen at multiple focal planes. When focused was set to the nerve fiber adjacent to the rim (Fig.3C) these structures were barely visible but as we moved the focal plane deeper, they began to appear (Fig.3D-E). When we focused down to the photoreceptor layer the signals of these circular structures started to become blurry again (Fig.3F), suggesting that they reside between the PR layer and GCL. They appeared similarly in another participant (SFig2C-F). Interestingly, we didn’t see these structures on all participants (SFig.2G-H).

**Figure 3.**
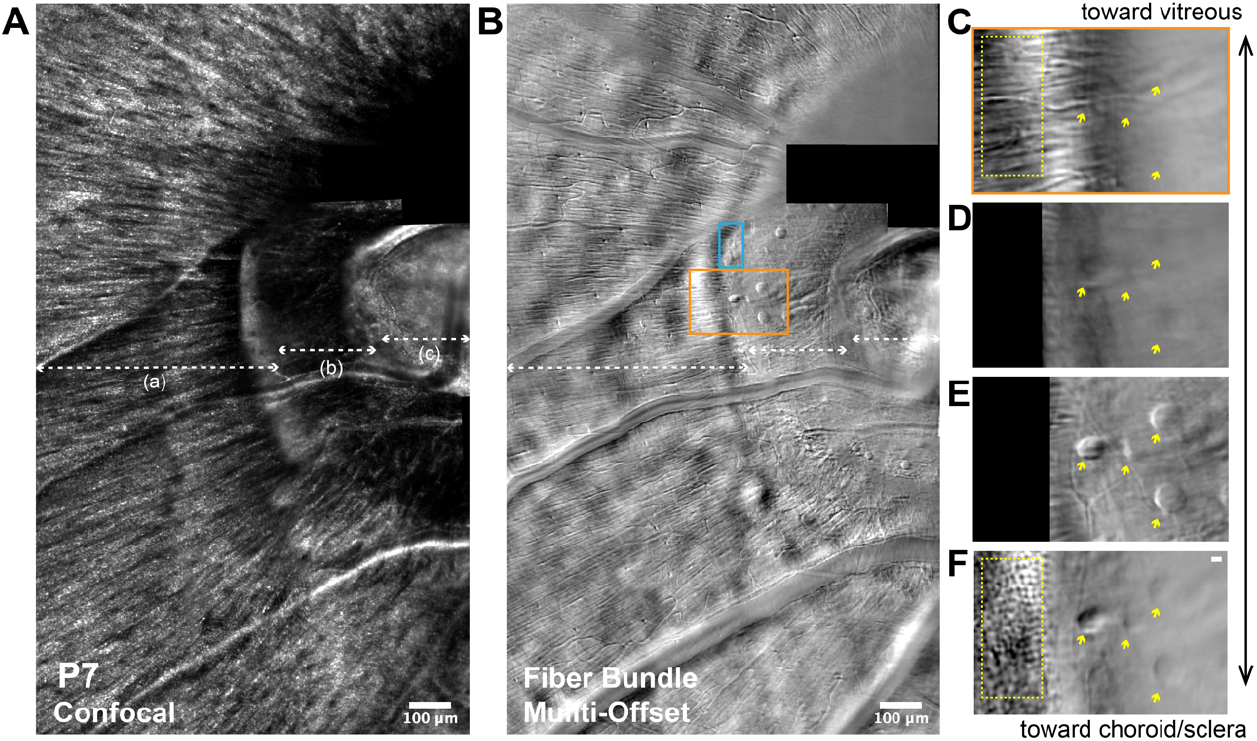
Optic Nerve Head region of a healthy participant (P7). 24-year-old healthy male participant. Montage of optic nerve head (ONH) region imaged with confocal **(A)** AOSLO and **(B)** FB multi-offset AOSLO. **(B)** three dashed lines with double arrowhead marked as (a), (b) and (c) referred to boundary where focal plane changes. **(C-F)** enlarged view of the region marked in orange in (B) acquired with different system focal planes. Double arrowheads indicate the relative depth of the images. Scale bar for is 10 μm and applied to image (C-F). yellow arrowhead marked the same cells with different sharpness when focusing on different depth of the retina. Yellow dashed boxes showing the nerve fiber and photoreceptor at corresponding focal plane.

**Figure 4.**
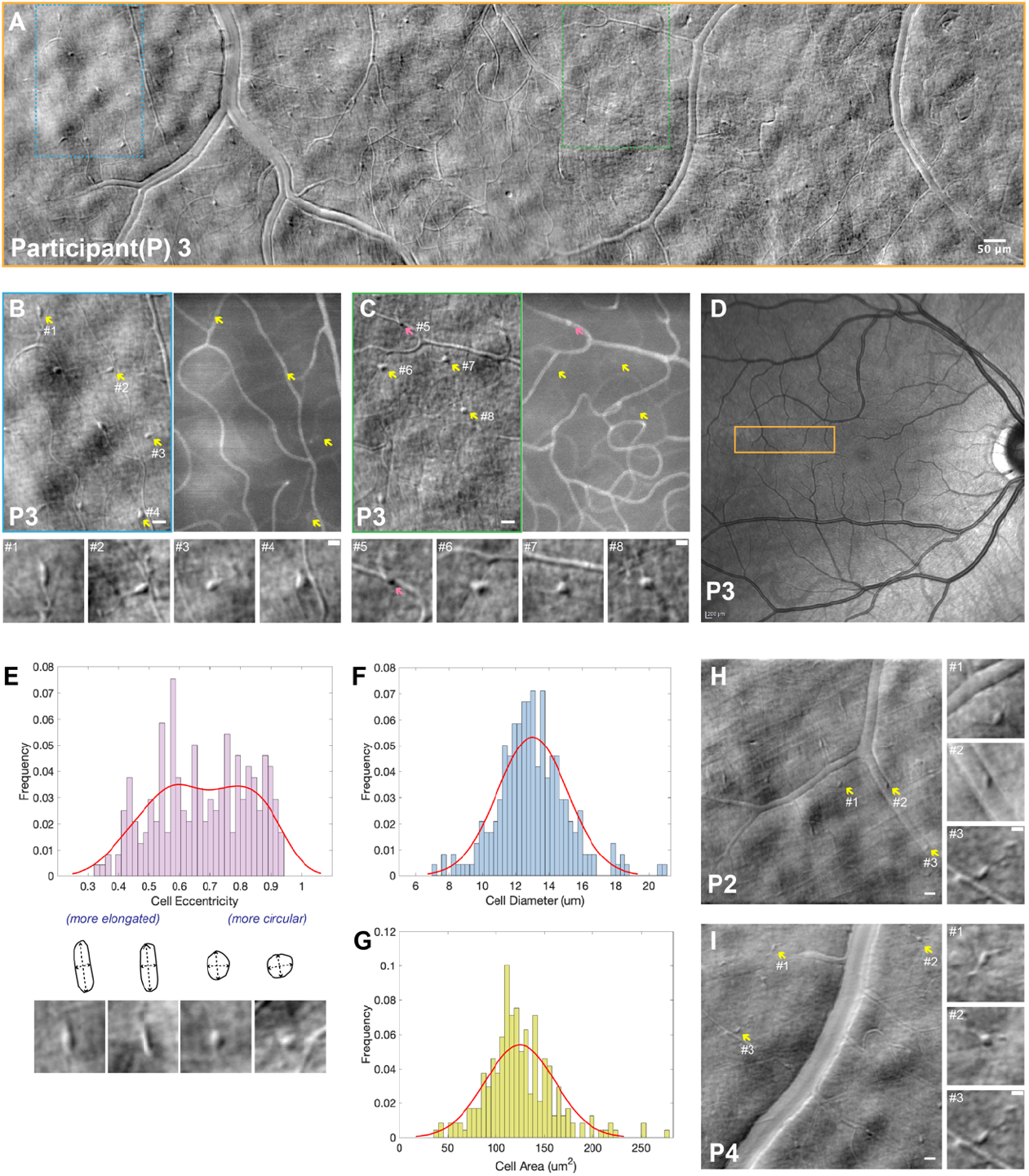
Microglia morphometry in healthy retinae. **(A)** retinal montage composed of 18 FB images. **(B-C)** images of blue and green region marked in larger montage in (A). Two larger images side by side on top are FB images and their perfusion maps, arrowheads pointed out example structures of different morphology with enlarged view shown below. Scale bars on larger images on top are 20 μm; Scale bars on smaller images on the bottom are 10 μm. **(D)** cSLO image with an orange box showed corresponding region of montage in (A). **(E-G)** histogram plot with distribution fit of all 239 cells from 6 participants: **(E)** histogram of cell eccentricity fit with a Kernel density estimation. Illustrations and example images below show microglia with higher and lower eccentricities, representing more circular cells and more elongated cells, respectively. **(F)** histogram of cell diameter fit with normal distribution (μ=13.02, σ=2.11). **(G)** histogram of cell area fit with normal distribution (μ=124.70, σ=35.88). **(H-I)** more examples of microglia with various morphologies detected on two additional participants. Larger images on the left are from one single acquisition (1.5°× 1.5° FOV) with enlarged views corresponds to yellow arrowheads showing zoomed in views of cell to the right. Scale bars are 20 μm for larger images; Scale bars for enlarged views are 10 μm.

### 3.2 Imaging presumed microglial cell in healthy retinas

Table 3 summaries the measurements made in the 6 normal eyes, where a total of 239 presumed microglia were detected. Fig.4A shows one of the large montages; dashed boxes are enlarged in Fig.4B-C. Microglia showed a spectrum of morphometries ranging from elongated, resembling ramified microglia, to a more circular amoeboid shape, resembling activated microglia. In some cases, microglial cell processes were clearly visible. To distinguish retinal vasculature from potential microglia processes, we compared the FB images to the simultaneously acquired perfusion maps generated through motion contrast. Yellow arrowheads in Fig.4(B and C) mark the position of seven cells that were seen in the FB images. Our hypothesis was that ramified microglia in the healthy retina would give little contrast in the perfusion maps, since they move little, while blood flow will contribute a strong motion contrast signal. Even if the microglia in the healthy retina moved, we hypothesized that they should still have lower contrast compared to the blood flow. Occasionally microglia-like structures were seen but when compared to the corresponding perfusion map, these showed high contrast and were co-localized with a vessel (pink arrowhead in Fig.4C). Owing to ambiguity as to whether this was a microglial cell or a diving vessel, we conservatively considered these structures to potentially be cross sections of vessels and excluded them in our quantification.

**Table 3.** Measurements of presumed retinal microglial cells detected in healthy retinae.

| P# | Eye | *Imaging region boundaries |  | imaging region (mm<sup>2</sup>) | microglial cell detected | Cell Density (/mm<sup>2</sup>) | Average cell area (μm<sup>2</sup>) | Average cell Diameter (μm) | Average cell Eccentricity | locations cell motility | cell monitored for motility | Speed (μm/sec) |  |  |
| --- | --- | --- | --- | --- | --- | --- | --- | --- | --- | --- | --- | --- | --- | --- |
|  |  | retinal field (number of images) |  |  |  |  |  |  |  |  |  | Average | Max | Min |
| 1 | OD | 7T ~ 12T; 2S ~ 6S | (20) | 1.73 | 65 | 37.5 | 125 | 13.26 | 0.62 | 3 | 12 | 0.042 | 0.106 | 0.020 |
| 2 | OD | 8T ~ 12T; 4S ~ 7S | (9) | 0.93 | 30 | 32.1 | 136 | 13.57 | 0.69 | 1 | 5 | 0.023 | 0.040 | 0.020 |
| 3 | OD | 2T ~ 12T; 0S ~ 3S | (18) | 1.41 | 61 | 43.3 | 134 | 13.54 | 0.68 | 3 | 16 | 0.007 | 0.016 | 0.002 |
| 4 | OD | 4T ~ 10T; 3S ~ 6S | (10) | 0.74 | 23 | 31.1 | 120 | 12.69 | 0.73 | 2 | 9 | 0.006 | 0.014 | 0.003 |
| 5 | OD | 2T ~ 11T; 0S ~ 2S | (18) | 1.28 | 40 | 31.2 | 106 | 11.78 | 0.74 | 1 | 2 | 0.008 | 0.009 | 0.007 |
| 6 | OS | 7T ~ 14T; 3I ~ 0I | (21) | 1.68 | 20 | 11.9 | 120 | 12.70 | 0.66 | 0 | N/A | N/A | N/A | N/A |
| summary |  | N/A (16) |  | 1.30 ± 0.4 | 40 ± 19 | 31 ± 11 | 123.5 ± 11 | 12.9 ± 0.7 | 0.7 ± 0.04 | total: 10 | total: 44 | avg: 0.017 | max: 0.106 | min: 0.003 |

\* Imaging region boundaries means the range of retinal eccentricity of the montage image formed with multiple FB images.

We measured cell eccentricity to describe their morphology and created a histogram plot for all cells detected (Fig.4E); a Kernel density estimation was fit to the plot that showed a double humped shape suggestive of potentially capturing cells in two different morphometries as defined by their eccentricity (i.e., circularity), a more elongated population centered around 0.58 and a more circular population peaking around 0.8. This agreed with our qualitative assessment that two main types of morphology were seen. The cell diameter of all 239 cells were plotted in a histogram (Fig.4F) and fit with a normal distribution (μ=13.02, σ=2.11). A histogram of cell area is plotted and fit with a normal distribution (μ=124.70, σ=35.88) in Fig.4G. Additional examples of microglia morphology from other participants is shown in Fig.4(H-I) and SFig.3. Microglia density was 31 cells/mm^2^, on average. However, we observed large variability in the density of cells across the 6 participants, ranging from 12–43 cells/mm^2^ (Table 3).

Microglia motility was measured at 10 different locations in 5 healthy eyes (Table 3). In healthy retinae, microglia moved extremely slowly, with an average speed of 0.02 μm/sec. Most healthy microglia were static and only a few were observed moving at higher speeds, of up to 0.11 μm /sec (Fig.5G). We plotted the cumulative distance against the observation time for 10 cells displaying a range of speeds from 3 different retinal locations from P1 in Fig.5(D). Here 10 consecutive images were generated across 5 minutes at an interval of 30 seconds. Here we show example images from 0, 3, and 4.5 minutes (Fig.5A-C). The purple line plot shown in Fig.5D has the highest speed of all cells monitored normal healthy eyes (corresponds to cell in purple box in Fig.5A). Supplementary Movie 2 shows the complete time-lapse animation from Fig.5A. It is also worth mentioning that cell speed between adjacent observation time points varies, indicating the uneven speed of cells that occasionally move in short bursts (e.g., cell plotted in purple between timepoints 2.5 and 3 in Fig.5D). Apart from changing their positions, we also noticed microglia changing shape to different degrees. This indicates that our microglia morphometry measurements should be taken to represent a snapshot in time and that parameters such as the eccentricity of cells can change dynamically over short timescales. The observation of morphological changes could also be indication of cells extending and retracting their processes (see yellow and light blue boxes in Fig.5E). Supplementary Vid 3-5 shows three more examples of cells changing shape over time.

**Figure 5.**
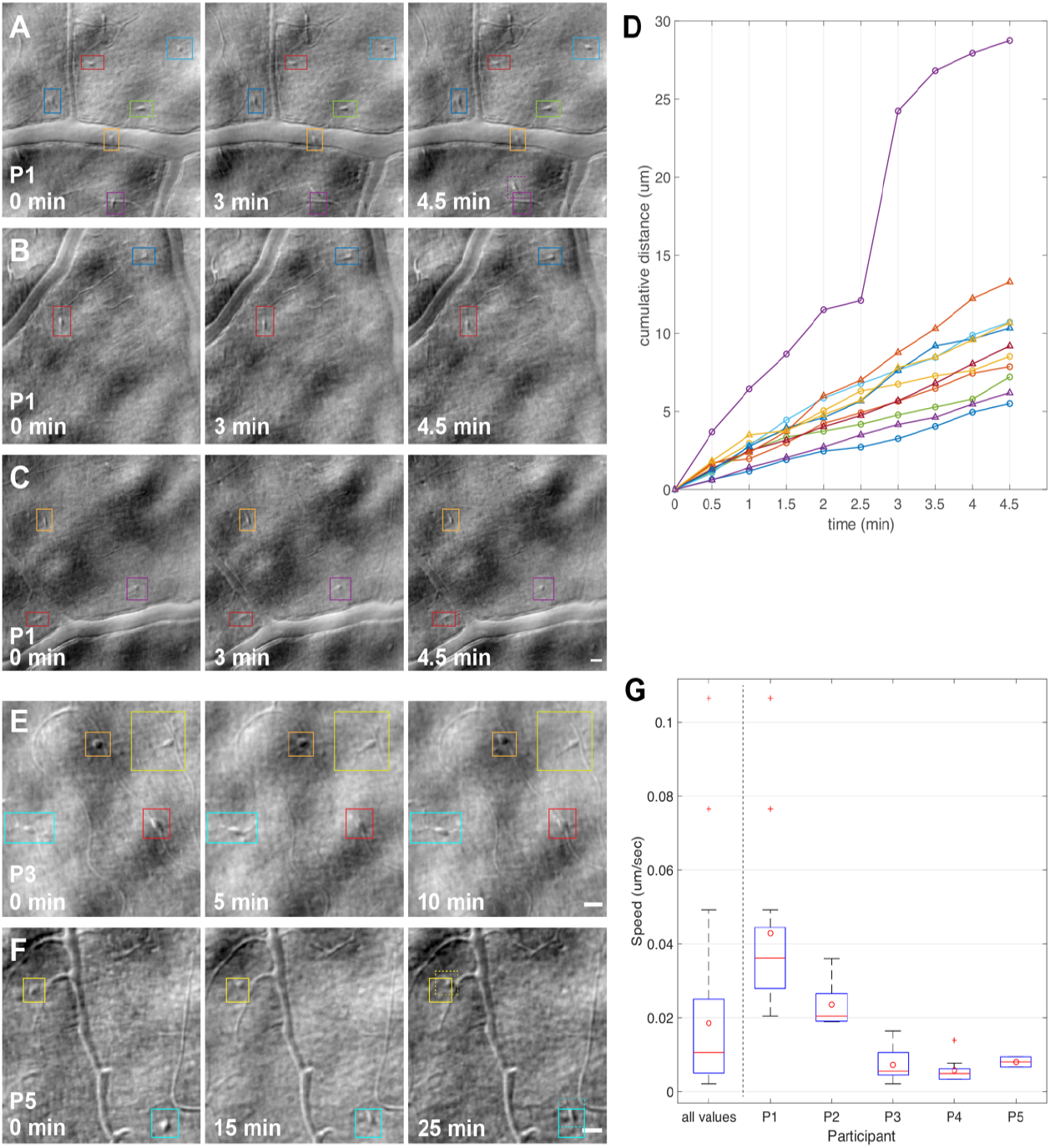
Motility of retinal microglial cells observed in healthy retinas. **(A-C)** 3 groups of images from 3 retinal locations of participant 1. For each location, 3 images from a 10-image sequence are shown, at 0min, 3min and 4.5min. Color boxes marked microglial cell positions and stays the same at each time point. When cell moves outside the initial solid box, we used dashed boxes to indicate its current position. **(D)** Plot of cumulative distance of 10 cells detected from (A-C) again the observation time point. **(E-F)** Two more examples of microglial cell motility in two other participants at three different observation time points. Color boxes highlight the cell positions and stay the same at all three images; dashed boxes denote cell that moves outside to initial solid boxes. **(G)** boxplot of microglial cell motility from all and each individual healthy participant imaged.

### 3.3 Monitoring cell dynamics in patients with retinal inflammation

At the first timepoint for imaging the ASPPC patient (Pt1) (Fig.6A) many presumed microglia and/or immune cells were observed in FB channels; careful comparison revealed some of these were also barely visible in the confocal channel (Fig.6B-C). Here we saw additional morphologies suggestive of macrophages. Some of these macrophage-like cells contained internal structures while others had more uniform contrast across their surface. Fig.6D-E shows examples of the cells on his first visit and in the two follow-up visits. At the initial visit during the most acute phase of the disease we observed a total of 32 cells in this field of view with an average cell diameter of 12.5 μm. We imaged this location multiple times on this first imaging session to assess cell motility. The intervals were 30 seconds apart for the first 3 minutes (6 images) and 30 seconds apart 15 minutes later (2 images). As described previously, to better capture the fastest moving cells, we used a 10-second integration time to evaluate fast moving cells within these sequences. We selected 10 cells with varying speeds across the range observed and highlight them in Fig.7(A-C). The complete time-lapse video is shown in Supplementary Vid 6 with white arrowheads denoting the positions of the fastest-moving cells. Due to limitations in graphical presentation of timelapse imagery we only show part of the observation time points in Fig.7A-C. Colored boxes are drawn in the same way as Fig.5. For the complete observation sequence, we extracted cell position values, re-drew the cell centroids, and connected the dots from the same cell to create the cell track plots shown in Fig.7D with the tracks overlaid on the first FB image. Longer tracks denotes cells that travelled longer distances. Some aggregated dots are zoomed-in and shown Fig.7E, to denote motions of cells that move only a small amount.

**Figure 6.**
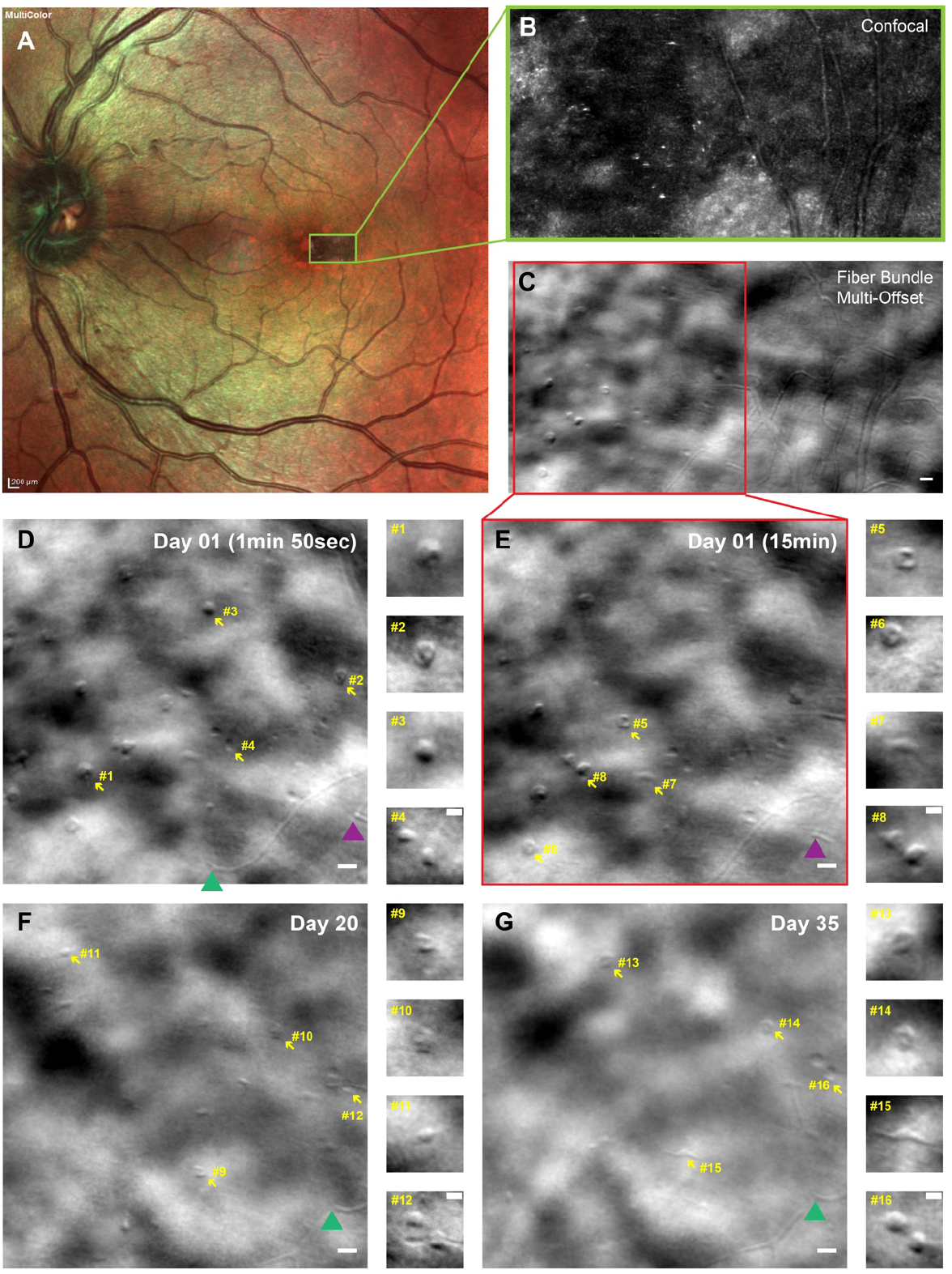
Left eye from patient number 1 with acute syphilitic posterior placoid chorioretinitis (ASPPC). Orientation of imaged retinal location and cells with different morphology. **(A)** Multicolor-mode fundus photo revealing granular dots within the foveal area; yellow box denoting region imaged. **(B-C)** confocal and FB images of affected region. Red box highlights region that we focused on for timelapse imaging and quantification. Scale bar is 20 μm and applies to B and C. **(D-G)** single FB images acquired at initial and follow-up imaging sessions. Yellow arrowheads denote cell with different morphology and was enlarged on the right. Green and purple triangle marked the same vessel landmarks at different imaging day and time for orientation. Scale bars on larger FB images are 20 μm; scale bars on smaller images on the right are 10 μm.

**Figure 7.**
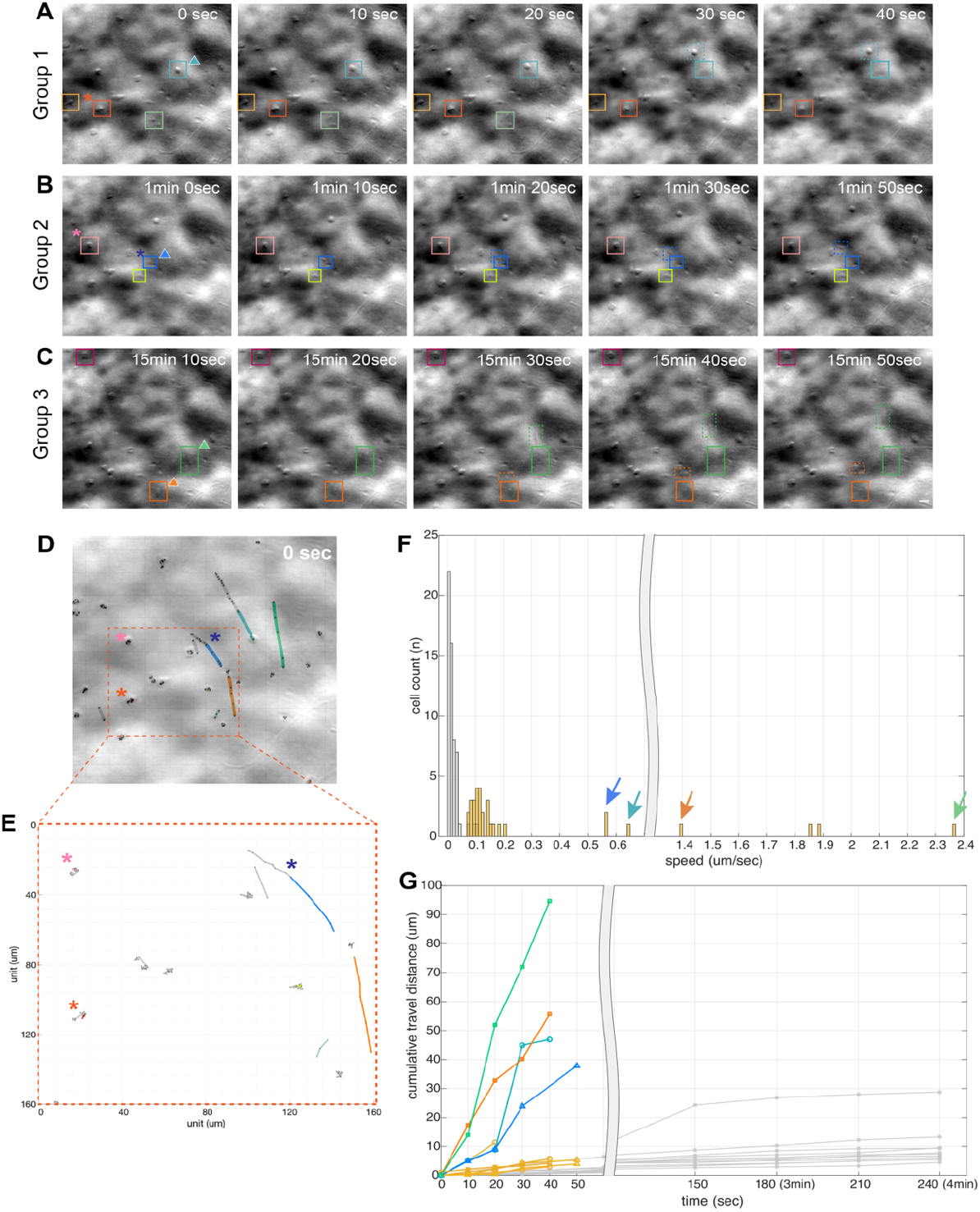
Microglial cell motility of patient number 1 (Pt1) with ASPPC. **(A-C)** 3 groups of cells selected from the entire observation showing cell position changing over time. Colored boxes marked cell location at different timepoint, and dashed boxes marked cell that moves outside initial boxes. Scale bar is 20 μm and applied to all A-C. **(D)** cell position track plot of cell detected on Pt1 and superimposed with first image showing the position of cells throughout entire observations. Colored part of line represents the images shown above and gray part represents cell position at observation time points that are not shown in this figure. **(E)** an enlarged view of cell track from (D) showing cells that moved locally and cells that travelled a long distance. **(F)** histogram plot of cell speed from Pt1 (yellow) and healthy participants (gray). Color coded arrowheads denote to cells marked with boxes in A-C and line plot in (G). **(G)** line plot of cumulative travel distance against observation time points showing the cell speed. Gray lines denote to microglial cells from healthy retinas, color coded line plots denote to cell speed corresponding to cell marked in A-C and F.

Cell motility was compared to normal eyes; Fig.7F shows a histogram of cell speed. Yellow bars denote cells detected in Pt1 while gray bars show the cells measured from healthy eyes. We selected the cells with highest speeds (marked with colored arrowheads) and plotted their cumulative distance in Fig.7G; gray line plots show the fastest microglia detected for healthy retinas. It should be noted that the fastest cells were not seen on all images because they rapidly moved outside the field of view. Compared to the slow motion of microglia seen in healthy retinas we observed substantially faster cells in this patient at this first timepoint, with speeds up to 2.4 μm/sec. Fewer cells were detected, and their average motility was reduced at the same location on follow-up imaging. This reduction in the number and speed of cells was correlated with improvements in the patient’s vision and his clinical exams and images. FBAOSLO images are compared across the different timepoints in Fig.8A-C. Orange boxes denote the same FOV through the 3 visits. Colored arrows stay at the same position relative to the orange boxes and are for orientation purposes. Histograms of cell speed for each timepoints are arrayed vertically in Fig.8D-F to appreciate both the reduction in the number of cells and their speed over time. SFig.4 shows comparison of clinical images and rtx1 images obtained on the 3 visits. Supplementary Vid 7-8 shows the complete time-lapse sequences for the second and third visits.

**Figure 8.**
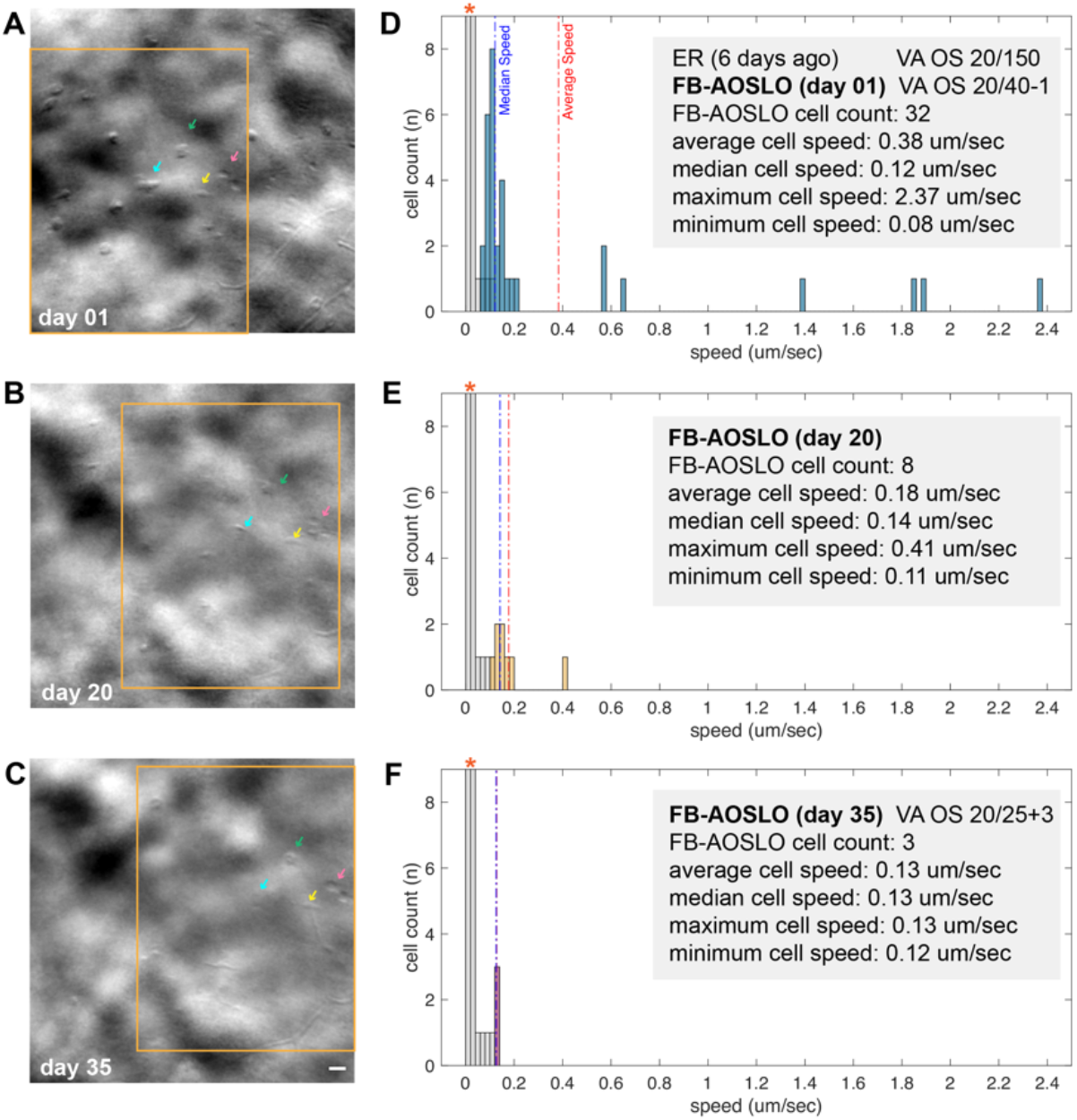
Comparison of Pt1 with ASPPC at 3 different FB-AOSLO imaging sessions. **(A-C**) FB images at different imaging day. Orange boxes showed the same retinal location. Colored arrowheads stayed at the same position relative to orange boxes and are for orientation purpose. Scale bar is 20 μm and applied to image A-C. (D-F) histogram of the speed of cell detected at corresponding imaging session. Gray bars stand for values from healthy retina, red start represents values that exceed axis limits. Colored bars stand for values from current patient. Dashed red lines and blue lines marked the average and median speed value for each imaging session. Notes on the gray background record key values at 3 different imaging sessions.

In the cases of Pt2 and Pt3, numerous microglial cells are seen as shown in Fig.9. These microglia have similar morphology and size compared to the microglia we detected in normal eyes but were substantially increased in number. Cell activity was reflected by cell morphology changes locally (Fig.9G-L) rather than by observation of them travelling over long distances, as was seen in patient 1. Cell activity (i.e., motility) was apparent over short intervals in an eye of a patient affected by a first episode of acute retinal vasculitis (Pt2) but absent in another patient (Pt3) affected by chronic retinal vasculitis that was moderately active at AOSLO imaging visit #1 and presumably quiescent at AOSLO imaging visit #2. For Pt3, the lack of cellular motility at visit #1 in suggests that microglia, at least at the location we imaged, were in a mild activated state because of mild macula edema and at visit #2 AOSLO images showed inactivated quiescent state. Conventional retinal imaging in Pt3 was not accurate enough to confirm uveitis activity. The presence of macula edema shown on OCT suggests a moderate level of persisting inflammation while on the second visit (6 months after intravitreal steroids injection) her uveitis was considered as presumably quiescent due to resolved macula edema. Her fluorescein angiography (FA) was persistently showing diffuse staining from the retinal vessels in both eyes due to chronic rupture of blood retina barrier, making the level of inflammation impossible to assess in that patient with >8 years of uveitis (Fig.10B-C) whereas FB AOSLO taken on two different days (Fig.10 D, G, J, and K) showed the different degree of cell infiltration that were not significant observed in clinical images. Comparing with Pt4 with chronic uveitis for over 20 months, she (Pt4) received intravitreal injection of steroids 6 weeks and 7 months ago (6 months sustained-release dexamethasone intravitreal and 36-month sustained-release fluocinolone acetonide, respectively) for macular edema before AOSLO imaging and it was resolved at the time of imaging and therefore chronic uveitis was quiescent and no microglial cells infiltration was observed.

**Figure 9.**
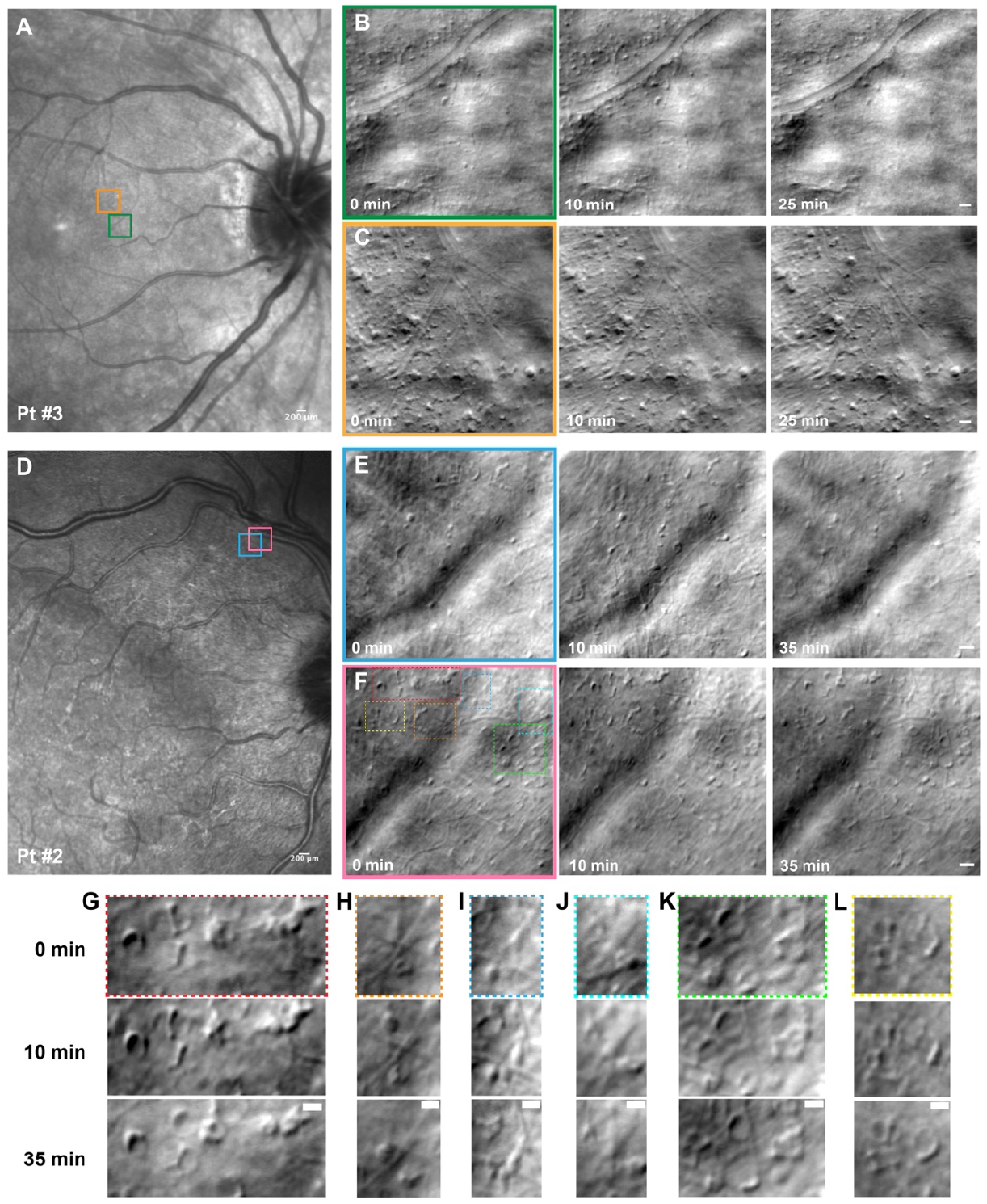
Comparison of FB images between patient with chronic (Pt 3) and active uveitis (Pt 2). **(A)** SLO images of Pt 3. **(B-C)** FB images of microglia imaged at the location indicated in the colored boxes in (A). **(D)** SLO images of Pt 2. **(E-F)** FB images of microglial cell imaged at the location indicated in the colored boxes in (D). **(G-L)** enlarged view of cells marked in F showing changes of cell morphology at three different time points. Scale bar for image B,C,E,F is 20 μm; Scale bar for image G-L is 10 μm.

**Figure 10.**
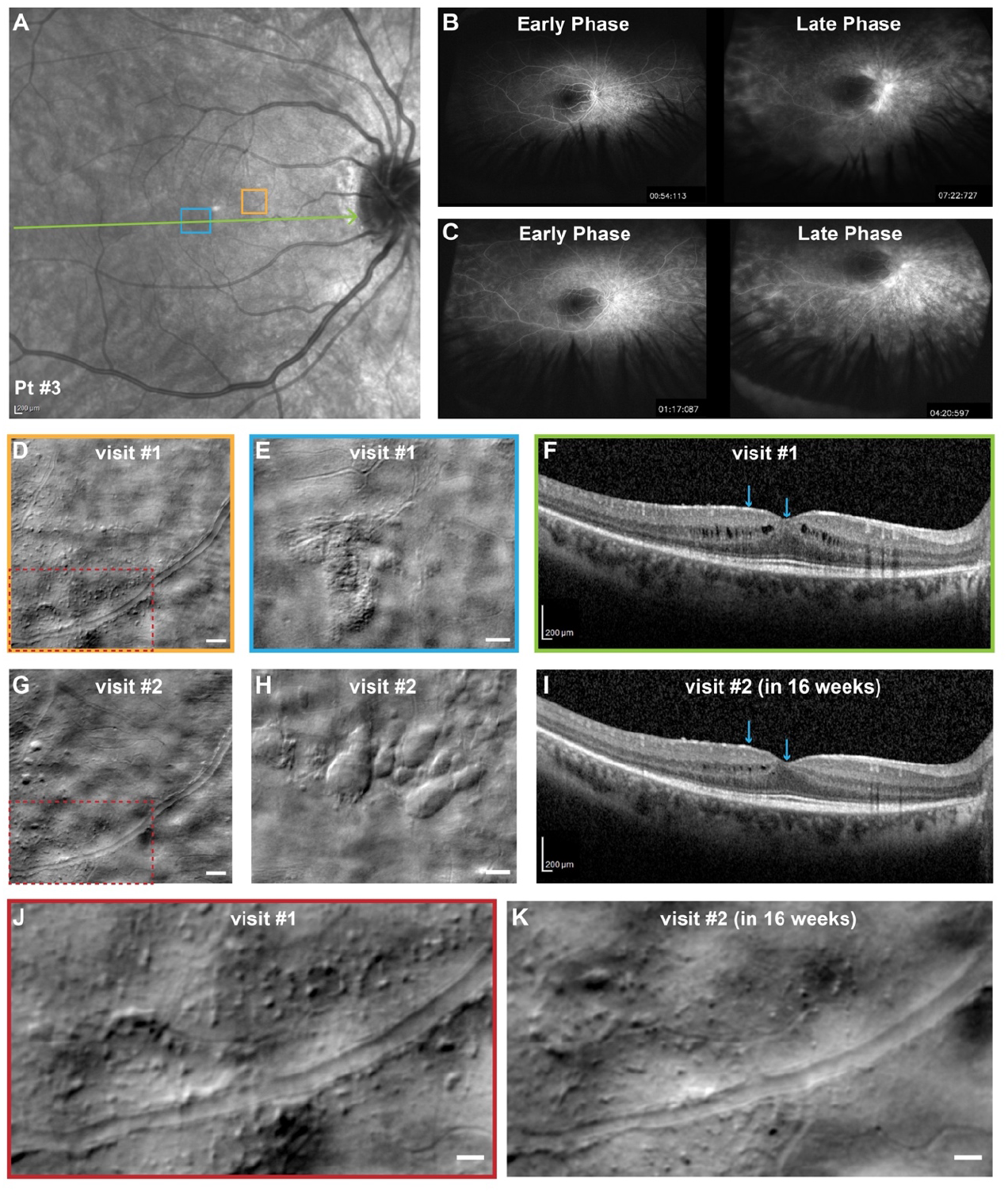
Images of Patient number 3 (Pt3). A 36-year-old female with chronic bilateral idiopathic posterior uveitis over 8 years. **(A)** SLO images of the patient. Green line showed OCT scanning slices. **(B)** FA at visit #1 showed chronic unchanged staining. This FA was done 3 months after periocular triamcinolone (steroids) injection, while on adalimumab since the last 5 months. **(C)** FA at visit #2 showed chronic unchanged staining. This FA was done 6 months after intravitreal steroids injection, while on adalimumab SQ since the last 12 months. **(D&G)** affected region with numerous microglial cell infiltrating. Scale bars are 50 μm **(J&K)** zoomed in view of (D) and (G), Scale bars are 20 μm. **(E&H)** the presence of macular edema. Scale bars are 50 μm. **(F&I)** OCT slices corresponding to green line in (A), blue arrowheads denote image boundary in E and H. Intraretinal macular edema was reduced, almost resolved in (I) (visit #2) as compared with (F) (visit #1).

## 4. Discussion

Fiber bundle AOSLO imaging of the human retinal ganglion cell layer reveals the fine structural details of microglia in healthy eyes and their limited motion over time in the absence of infection or inflammation. This enabled visualization of microglia over larger areas and imaging of the ONH region. Near the ONH we detected for the first time many cell-like structures that may represent astrocytes or other macroglia. In patients, we observed the activity of presumed microglia and macrophage-like cells in a spectrum of disease states including acute syphilitic infection where we saw many macrophage-like cells and observed rapid motility. In patients with active or chronic uveitis, the presence of microglia motility was observed in an active case, but motility was absent in the case of a chronic uveitis patient receiving immunomodulatory treatment. These observations in patients with uveitis and active ocular infection demonstrate the potential for future imaging studies using this technique and suggest that there is substantial clinical potential for using FB-AOSLO clinically to study human disease.

The major advantage of fiber bundle detection is the simultaneous acquisition of all offset positions with reduced imaging duration, and identical AO correction and field of view. Mozaffari et al.^28^ first showed the feasibility of such an approach and we implemented it here using a custom fiber-bundle that mimicked our previously shown improved multi-offset detection pattern. Comparing the images acquired from sequential detection and FB detection, no visible difference of image quality was seen, and their normalized power spectral density showed FB detection can improve the image quality slightly. When testing different exposure times, no visible differences in image quality were seen from 20–60 seconds, and this was also validated from plotting their normalized energy profile. This demonstrated that there was no need to extend the single acquisition time beyond 20 seconds when imaging healthy eyes. Here we used 30 seconds rather than 20 seconds based on the consideration of older participants and patients. However, we showed here that high contrast images of a patient with a severe ocular infection (Pt1) could be generated using 10 seconds of data to increase the temporal sensitivity for motility quantification, so shorter intervals are certainly feasible. One aspect that could be improved to increase the signal at the detectors and potentially shorten the interval would be to provide some collimation of the light at the detection end as we calculated that given the NA of the fibers used that we were slightly overfilling the PMT photocathode.

Another advantage of FB imaging is the simplified optical implementation that removes the need to form multiple retinal conjugate focal planes. This simplifies the alignment over approaches such as split- and quad-detection imaging, where multiple optical elements are needed to divide up the confocal and non-confocal light. This allowed us to use participant fixation along with our control of the imaging field of view using the steering mirror to generate images of the optic nerve head region that was previously inaccessible in our sequential multi-offset setup. We did notice however that certain elderly patients with intraocular lens (IOL) were challenging to image, especially near the ONH. We found this begins to influence the image quality when exceeding 14 degrees away from the fovea. Considering the population of people with glaucoma and IOL, this should be further investigated to overcome this limitation.

An important consideration is that we found the focus slightly varied as we moved across the retina to generate large montages, likely due to changes in retinal thickness and keeping the focal plane on the GCL was more challenging than for the outer retina, where the cones provide a strong indicator of the focal plane. It should be noted that keeping the focal plane constant is important for making larger montages at all retinal regions and especially when imaging areas such as the ONH, where the retinal thickness is rapidly changing, this deserves careful attention and thoughtful interpretation. When imaging the circular structures we detected on the optic rim, we imaged these structures at multiple focal planes but as mentioned previously this was not as systematic as it can be when imaging other areas of the retina. We were able to estimate the relative position of the cell-like structures seen in this area to show that they exist at several focal planes from the RGC layer to the photoreceptor layer. Based on their size and location at multiple stratifications, we hypothesize that these structures may be the somas of astrocytes, but they could also represent other cell classes so further studies are needed to evaluate these structures. Nevertheless, these first glimpses of putative astrocytes and the high contrast images obtained of the ONH region suggest there is potential for FB imaging in glaucoma and other conditions where cellular level imaging of the ONH can provide insight into ocular disease.

We were inspired to monitor presumed microglial cells using FB-AOSLO based on our previous observations using sequential multi-offset where we saw cellular structures that changed their location over time^8^. These structures were detected when we focused on the retinal ganglion cell layer. As described in the methods section, we use the real-time confocal channel and offset channel images during acquisition to help us determine the focal plane. Since we used the visibility of the nerve fiber bundle as a reference and then carefully moved to a slightly deeper layer to target the GCL, our contention is that the cells we have imaged reside within that layer. However, due to the relatively poor axial resolution of confocal AOSLO compared to other modalities such as AOOCT and uncertainties about the axial resolution of FB-AOSLO we cannot be completely certain that we are not also detecting cells that reside on the surface of the RNFL. Therefore, future work is needed to compare the cells we detected here to those that were detected previously by others, such as the vitreous cortex hyalocytes that were detected recently by Migacz et al.^9^. Since those images show no trace of the RNFL we suspect that they must have been taken at a more vitread focal plane than we imaged here. However, since those cells can also be detected using commercial OCT, we plan in future investigations to compare the cells we can detect with our FB-AOSLO setup to those seen on OCT to better understand the layer at which the cells we are imaging reside. We also plan to evaluate the efficacy of FB-AOSLO for detecting microglia at other retinal layers, such as the plexiform layers where they are known to be most numerous.

With FB-AOSLO, we are now able to acquire multiple images across relatively short intervals to form time-lapse sequences that visualize microglia in motion. Time-lapse sequences of 10 images can be generated in as short as 5 minutes with FB-AOSLO, whereas using the sequential approach it could take half an hour or more. Such long imaging duration was insufficient for capturing fast moving cells and can potentially make the cells appear elongated artificially due to fusion of data across aperture positions with the cell at different locations. This prevented accurate detection and quantification of cell morphology and motion with sequential multi-offset imaging. With FB-AOSLO, we showed that microglia cells in healthy retinae have a spectrum of morphologies including rounded cells and elongated cells with visible processes. This is consistent with previous work from histology that has shown that microglia in the adult retina are seen in two types, ramified parenchymal microglia having characteristics of dendritic antigen-presenting cells and microglia around blood vessels similar to macrophages or cells of the mononuclear phagocyte series^35,36^. However, there are some limitations in our ability to detect the processes of cells, lending some uncertainty to how the cell morphology might vary over time. For example, for the cells that appeared more circular, there is the possibility that we were only capturing part of their somas that appeared circular, this is also reflected by the fact that we were not able to visualize all the processes of these cells. Future work is needed to improve the image processing for better detection of cellular processes. The use of filters to enhance particularly oriented spatial frequencies may be efficacious for this purpose, as was shown previously with the emboss-filter approach used by Migacz et al.^9^.

Imaging microglia in vivo in humans with FB-AOSLO has several limitations compared to in vitro preparations in animals or of human tissue samples that can be validated with histology and can capture the complete distribution and morphologies. However, the ability of in vivo imaging of human retinal microglia and macrophage-like cells has the advantage of monitoring their changes over short timescales and longitudinally, offering new opportunities for the evaluation of retinal diseases with active retinal changes, such as retinal infection and inflammation. This could reveal the severity of the disease and at the same time indicate the level of activity of retinal inflammation in a quantitative way. Though AOSLO imaging could sometimes be ineffective due to media opacities such as vitritis or cataract, it offers the unique possibility of quantifying the detailed dynamic features of individual cells in posterior uveitis when the media is clear enough for imaging. Considering future use of FB-AOSLO as a tool for studying uveitis, improvements are needed to replace the manual procedures we used here for quantification that would be insufficient for larger scale studies with automated methods for cell detection and tracking.

Non-confocal AOSLO imaging of mice has allowed label-free imaging of microglia and their motility^37,38^. Some of these investigations in mice have used fluorescent markers to identify resident microglia and differentiate them from infiltrating immune cells as well as image retinal inflammation in an endotoxin induced uveitis model^39^ where heterogenous immune cells, neutrophil and monocyte populations and their motility were imaged. They also showed neutrophils rolling along the venular endothelium and infiltrating monocytes and macrophages present both in vessels and extravasated into retinal tissue^39^. One of the limitations of human imaging is that we do not have the flexibility to selectively label cells, so we lack the capability to differentiate activated resident microglia from infiltrating macrophages. However, many of the images published from these previous mouse studies look strikingly similar to what we observed here in patients with uveitis, particularly the macrophage-like cells containing granular internal structures that we saw in a case of ASPPC (Pt1).

In the current study, cell motion differed markedly between active uveitis and healthy eyes without inflammation. Cells moved slowly in normal eyes (0.02 μm/sec on average, 0.1 μm /sec maximum), but their motility was increased in the eye with active inflammation secondary to infection (Pt1, ocular syphilis) and could move at speed up to 2 orders of magnitude greater than the normal average (up to 2.37 μm/sec). When observed after 3 weeks (visit #2) and 5 weeks (visit #3) of appropriate antibiotics treatment (penicillin IM), decreases in the quantity and motility of cells were correlated with improvements to vision and other structural and systemic biomarkers. This demonstrates the potential for FB AOSLO to evaluate treatment response at a cellular level in the living human eye. Careful comparison of the FB AOSLO images acquired from 3 different visits in this patient (Pt1) suggests that we were able to detect some of the same cells at all 3 visits. Colored arrowheads in Fig.8A-C denote the similar appearance, especially when comparing to the adjacent vessel landmarks. However, considering these images were acquired on different days, it is impossible to validate that these are the exact same cells.

To our knowledge, this is the first time that microglia and their motility have been observed during active infections or inflammations in the living human eye. Current commonly used clinical imaging modalities for managing posterior uveitis and particularly vasculitis include fluorescein angiography (FA), fundus autofluorescence imaging (FAF), and Optical Coherence Tomography (OCT)^23,24^. Only a few AO imaging case-reports and small case series have studied the photoreceptor layer and vasculature changes in uveitis but microglia have not been studied^25–27^. Using FB-AOSLO, we observed in one eye with infectious uveitis (i.e., ASPCC) the presence macrophage-like cells containing granular internal structures in GCL associated with chorioretinitis. The contrast was striking in motion between the cells observed in active uveitis and healthy eyes without inflammation. Cells moved slowly in normal eyes, but their motility was increased in eyes with active infection and inflammation. Uveitis treatment involves therapeutic immunomodulatory strategies. Those therapies may be categorized into those that modulate the general activation state of microglia, like corticosteroids and those that modulate specific molecular pathways through which microglia exert pathologic effects.^2^ Local retinal delivery of corticosteroids has demonstrated significant suppressive effects on microglial activation^40^. In our study, imaging by FB-AOSLO confirmed those findings showing only rare microglial cells in the GCL of a patient with chronic posterior uveitis and recent resolved macular edema two months after intravitreal steroids injection. This suggests that FB-AOSLO may be useful for monitoring the efficacy of immunomodulatory therapies in uveitis especially in case of chronic uveitis when the FA cannot ascertain the level of uveitis activity or quiescence because of chronic retinal staining.

Objective measures of intraocular inflammation are still a matter of current research for posterior uveitis. The amount of leakage in retinal vasculitis have been studied recently with development of automated leakage analysis of wide field fluorescein angiogram in uveitis patients. Those studies focus on automated processing techniques for standardization and normalization of fluorescein angiography images in patients with uveitis^41^. For retinal and choroidal lesions, autofluorescence and en face fundus images are now used in clinical trials to assess the level of inflammation. Other markers of inflammation are used in clinical practice, including macular edema, cells in vitreous (diagnosed clinically or by OCT), and clinical vascular sheathing. Better grading is essential for the development of disease modifying strategies to assess the efficacy of treatments. The development of clinical imaging tools for in vivo visualization of inflammation within the retina is currently needed to provide objective and quantitative proof of active versus quiescent ocular inflammation. FB-AOSLO imaging of the retinas of patients under immunomodulatory treatments for study of immune cells and their dynamics may allow for optimal understanding of immunological pathways in various human diseases and complement the existing clinical toolkit in the treatment of uveitis.

## 5. Conclusion

1. FB-AOSLO simplifies the optical design, improves the imaging efficiency, and extends the capabilities of phase imaging in AOSLO
2. FB-AOSLO permits the fine-scale structure of the GCL, including microglia and their motility, to be detected and quantified in normal eyes
3. FB-AOSLO reveals putative astrocytes in the living eye of humans for the first time
4. FB-AOSLO allowed the morphology and dynamics of microglia and macrophages to be quantified in the living eye of patients with uveitis and active infections for the first time
5. FB-AOSLO offers promise as a potential tool to evaluate disease status (i.e., active or quiescent) and response to immunomodulatory treatment in uveitis both clinically and in future trials.

## Supporting information

Microglial cell motility observed in healthy retina (participant 1) with random intervals.

Microglial cell motility observed in healthy retina (participant 1) with 30second intervals and corresponds to Fig.5A. White arrowhead tracks the posi

Microglial cell motility observed in healthy retina (participant 2). White arrowhead tracks the cell that changes morphology locally.

Microglial cell motility observed in healthy retina (participant 3) and corresponds to Fig.5E. White arrowhead tracks the cell that changes morphology

Microglial cell motility observed in healthy retina (participant 5) and corresponds to Fig.5F.

immune cell motility observed in patient 1 with ASPPC on the first visit. Solid cyan boxes on top marked the region of time-lapsed below. Dashed cyan

immune cell motility observed in patient 1 with ASPPC on the second visit. Solid cyan boxes on top marked the region of time-lapsed below. Dashed cyan

immune cell motility observed in patient 1 with ASPPC on the third visit. Solid cyan boxes on top marked the region of time-lapsed below. Dashed cyan

Microglial cell motility observed in patient 3 with chronic uveitis. Microglial cells barely moved. Correspond to Fig.9C.

Microglial cell motility observed in patient 2 with active uveitis. Microglial cells have relatively more activity by changing its morphology and posi

## Support

This work was supported by grants from Foundation Fighting Blindness (PPA-0819-0772-INSERM) and the BrightFocus Foundation (G2017082), departmental startup funds from the University of Pittsburgh to Ethan A. Rossi, NIH CORE Grant P30 EY08098 to the University of Pittsburgh Department of Ophthalmology, the Eye and Ear Foundation of Pittsburgh, and from an unrestricted grant from Research to Prevent Blindness, New York, N.Y. USA.

This work was also supported by Xiangya School of Medicine, CSU visiting scholar program.

## Acknowledgments

The authors would like to thank Austin Roorda for sharing his AOSLO software with us.

## Disclosures

Ethan A. Rossi has patents on some aspects of the technologies used to carry out these experiments (US10,772,496; US10,123,697; & US10,092,181). All other authors declare that they have no conflicts of interest related to this article.

## Data availability

Data underlying the results presented in this paper are not publicly available at this time but may be obtained from the authors upon reasonable request.

## Supplementary Materials

**SFigure 1.**
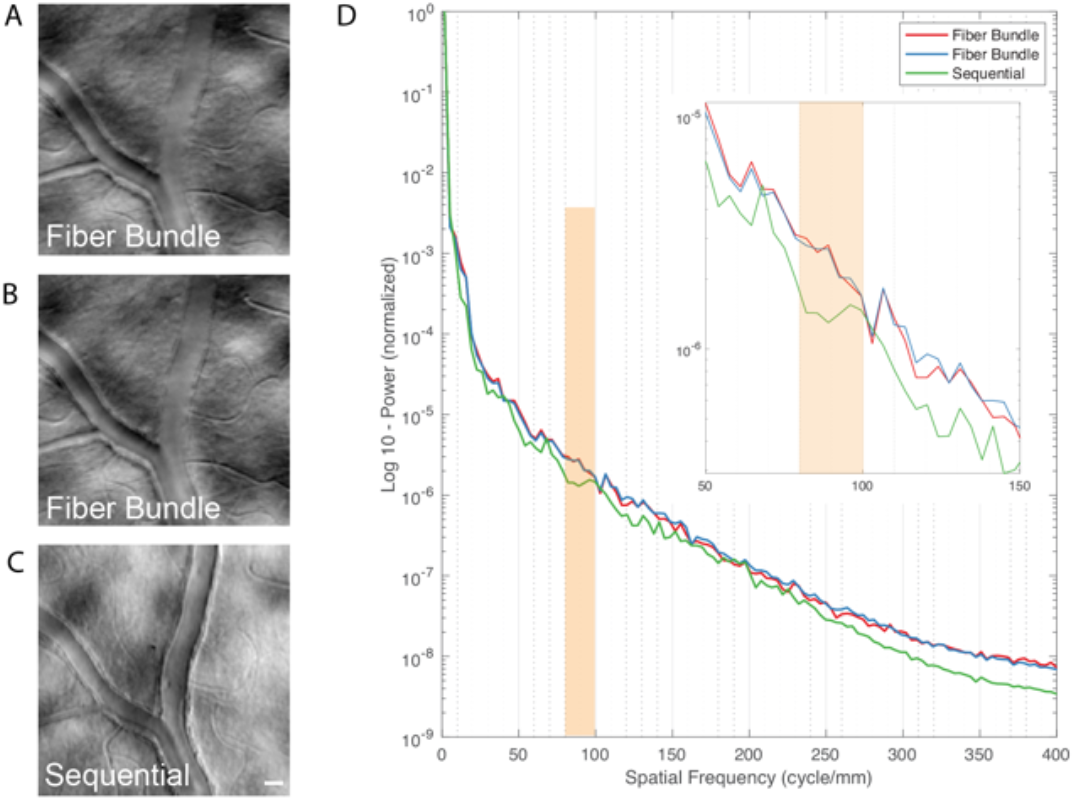
Comparing image quality between fiber bundle and sequential multi-offset imaging of participant 6. (A-C) images of 2 different depth of focus for fiber bundle and 1 for sequential detection. All three images have exposure time 30sec/aperture. Scale bar is 20 μm and applied to image and applied to image A-C. **(D)** normalized radially averaged PSD log plot of images (A-C). Orange box highlights spatial frequencies of interest. Insert plot enlarged the plot for spatial frequency ranging 50-150 cycle/mm.

**SFigure 2.**
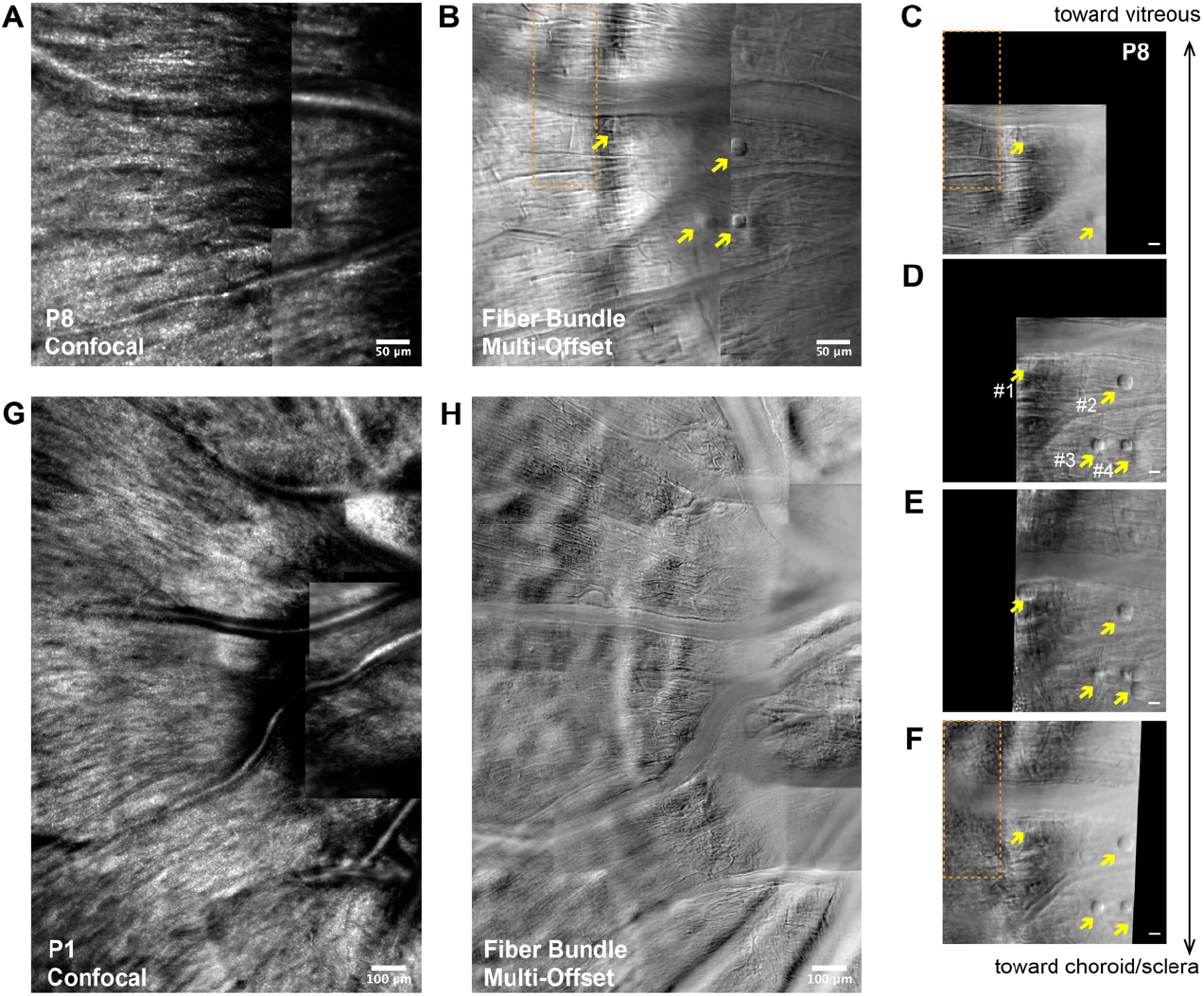
Optic Nerve Head region of 2 healthy participants. **(A-F)** 24-year-old healthy male participant. Montage of optic nerve head (ONH) region imaged with confocal **(A)** AOSLO and **(B)** FB multi-offset AOSLO. **(C-F)** enlarged view of cellular structures seen in (B) acquired with different system focal planes. Double arrowheads indicate the depth. Scale bar for is 10 μm and applied to image (C-F). yellow arrowhead marked the same cells with different sharpness when focusing on different depth of the retina. Orange boxes showing the nerve fiber and photoreceptor at corresponding focal plane. (G-H) 26-year-old healthy female participant. Montage of optic nerve head (ONH) region imaged with confocal **(G)** AOSLO and **(H)** FB multi-offset AOSLO.

**SFigure 3.**
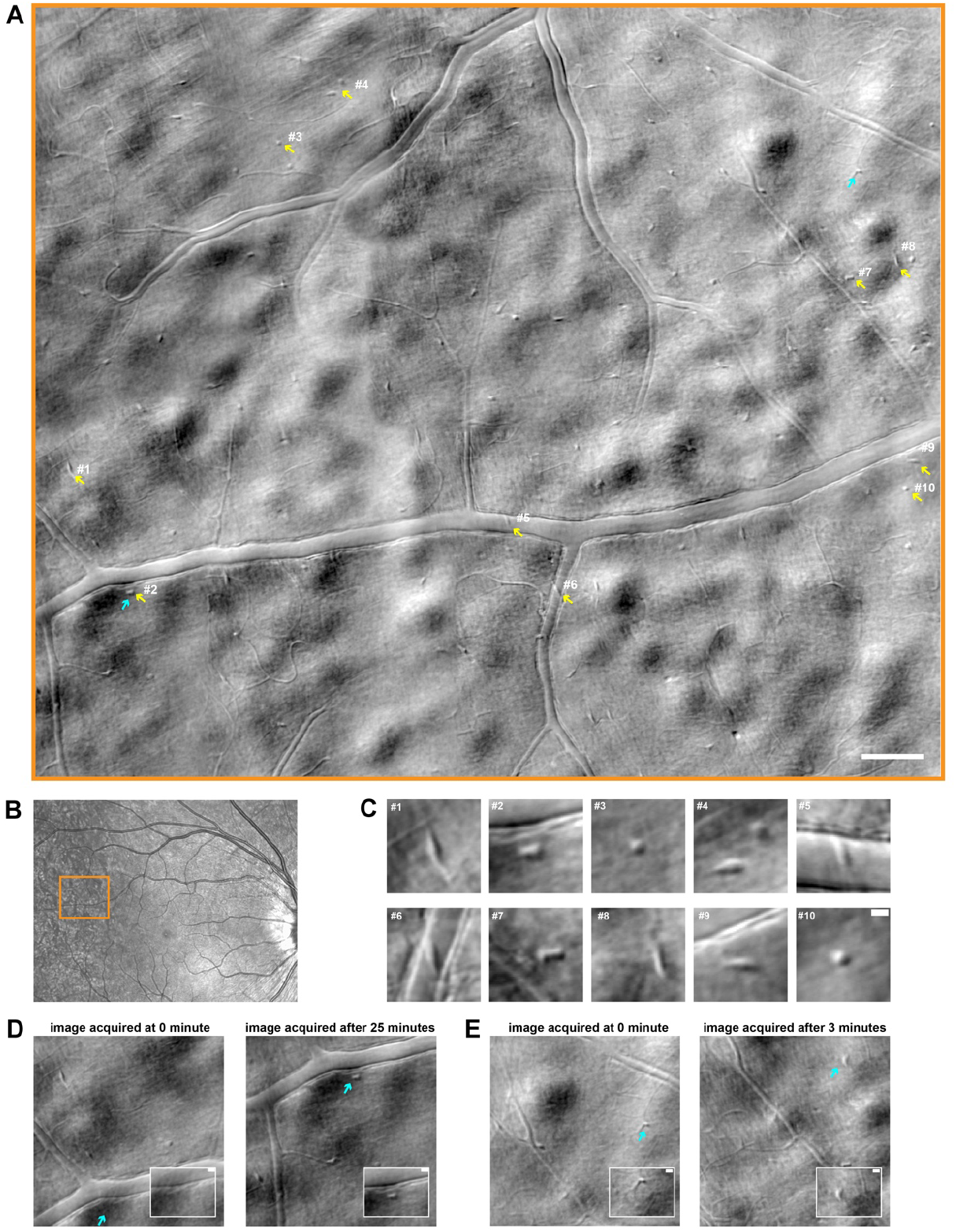
retinal microglial cells observed in healthy retina (Participant 1). **(A)** retinal montage composed of 20 FB images. **(B)** cSLO image with an orange box showed corresponding region of montage in (A). **(C)** a group of images showed enlarged view of microglial cells with different eccentricity marked in A. Scale bar is 10 μm and applied to all small figures in B. (D-E) showed two pairs of images shared mutual retinal location. Light blue arrowhead pointed at cells that appeared or changed morphology between two images. Image acquisition interval are labeled on the top of the image. Scale bar in the small insert figure is 10 μm.

**SFigure 4.**
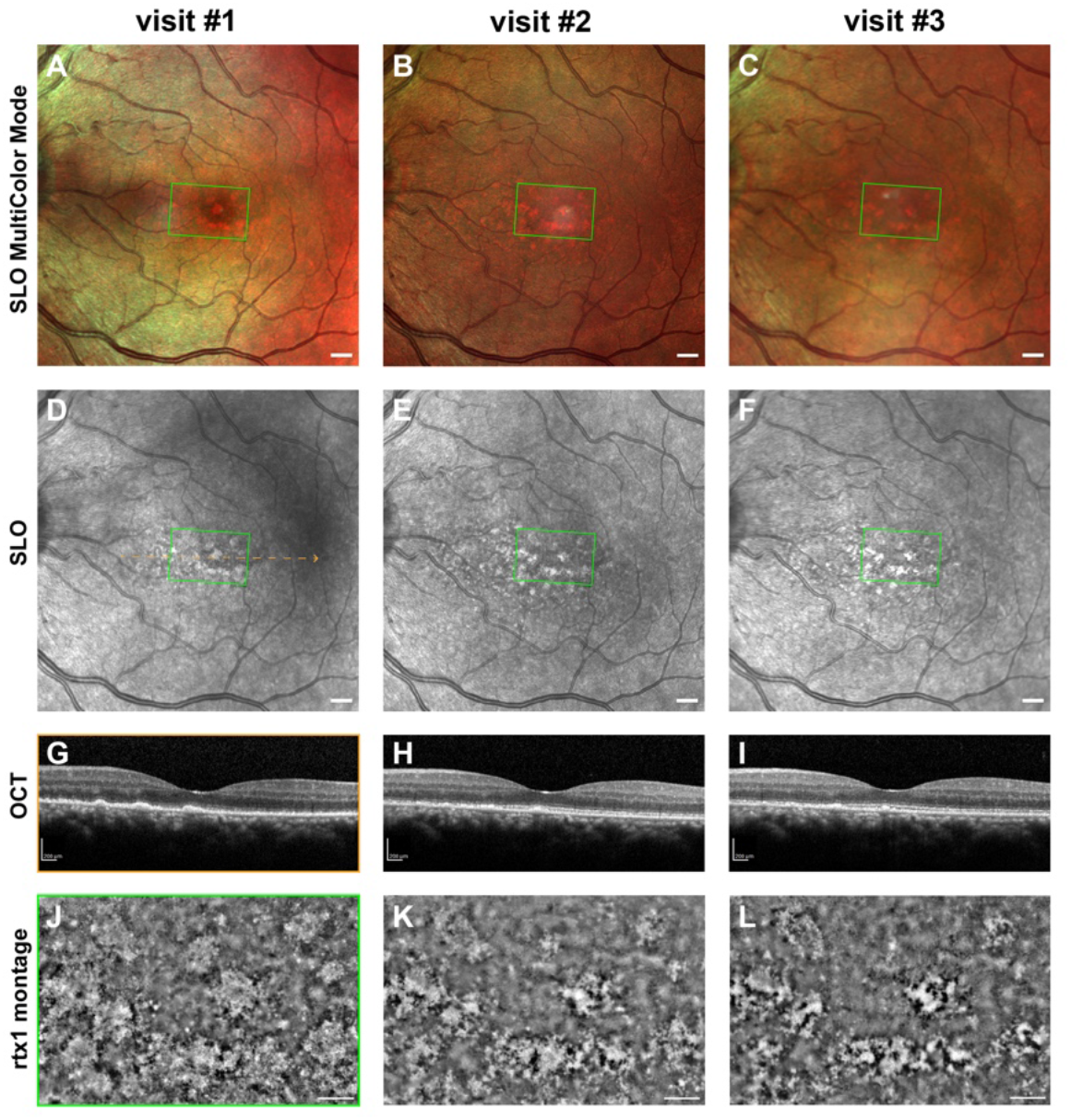
Clinical images and rtx1 retinal images of patient 1 with ASPPC at three different visits. **(A-C)** Multi-color mode fundus image. **(D-F)** grayscale cSLO images. **(G-I)** OCT slices of the retina marked with orange dashed line in D. (J-L) rtx1 retinal montage images of region marked in green boxes in A-F. (A,D,G,and J) are acquired on initial visit. Multi-color and grayscale SLO are acquired when patients admitted to ER 6 days prior to being imaged with AOSLO. (B,E,H,K) are acquired on the second visit. (C,F,I,) are acquired 2 days prior to the third AOSLO imaging due to patient’s personal arrangements, (L) is acquired on the third visits

**Supplementary Video.1** Microglial cell motility observed in healthy retina (participant 1) with random intervals.

**Supplementary Video.2** Microglial cell motility observed in healthy retina (participant 1) with 30-second intervals and corresponds to Fig.5A. White arrowhead tracks the positions of cell that moves the fastest of all.

**Supplementary Video.3** Microglial cell motility observed in healthy retina (participant 2). White arrowhead tracks the cell that changes morphology locally.

**Supplementary Video.4** Microglial cell motility observed in healthy retina (participant 3) and corresponds to Fig.5E. White arrowhead tracks the cell that changes morphology locally.

**Supplementary Video.5** Microglial cell motility observed in healthy retina (participant 5) and corresponds to Fig.5F.

**Supplementary Video.6** immune cell motility observed in patient 1 with ASPPC on the first visit. Solid cyan boxes on top marked the region of time-lapsed below. Dashed cyan boxes marked the region of time-lapsed for other imaging session. Dashed orange boxes denote the same retinal location across three different imaging sessions. Colored arrows in the time-lapse gif highlights fast-moving cells. Correspond to Fig.7 and Fig.8A.

**Supplementary Video.7** immune cell motility observed in patient 1 with ASPPC on the second visit. Solid cyan boxes on top marked the region of time-lapsed below. Dashed cyan boxes marked the region of time-lapsed for other imaging session. Dashed orange boxes denote the same retinal location across three different imaging sessions. Correspond to Fig.8B.

**Supplementary Video.8** immune cell motility observed in patient 1 with ASPPC on the third visit. Solid cyan boxes on top marked the region of time-lapsed below. Dashed cyan boxes marked the region of time-lapsed for other imaging session. Dashed orange boxes denote the same retinal location across three different imaging sessions. Correspond to Fig.8C

**Supplementary Video.9** Microglial cell motility observed in patient 3 with chronic uveitis. Microglial cells barely moved. Correspond to Fig.9C.

**Supplementary Video.10** Microglial cell motility observed in patient 2 with active uveitis. Microglial cells have relatively more activity by changing its morphology and positions locally. Correspond to Fig.9F.

## Notes

### Competing Interest Statement

The authors have declared no competing interest.

