## Supplementary figures and images for "Imaging the structure and dynamic activity of retinal microglia and macrophage-like cells in the living human eye"

### immune cell motility observed in patient 1 with ASPPC on the first visit. Solid cyan boxes on top marked the region of time-lapsed below. Dashed cyan

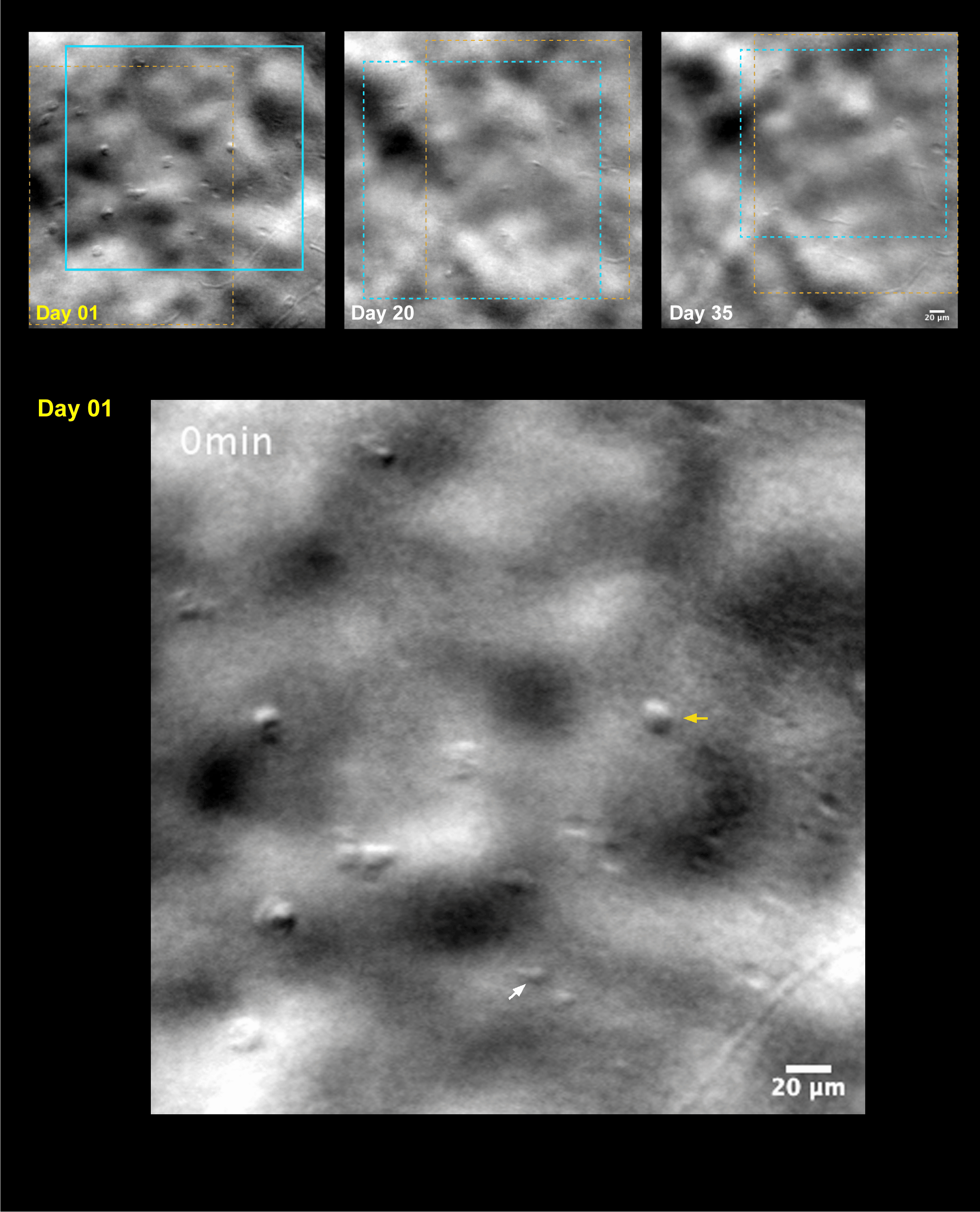

### immune cell motility observed in patient 1 with ASPPC on the second visit. Solid cyan boxes on top marked the region of time-lapsed below. Dashed cyan

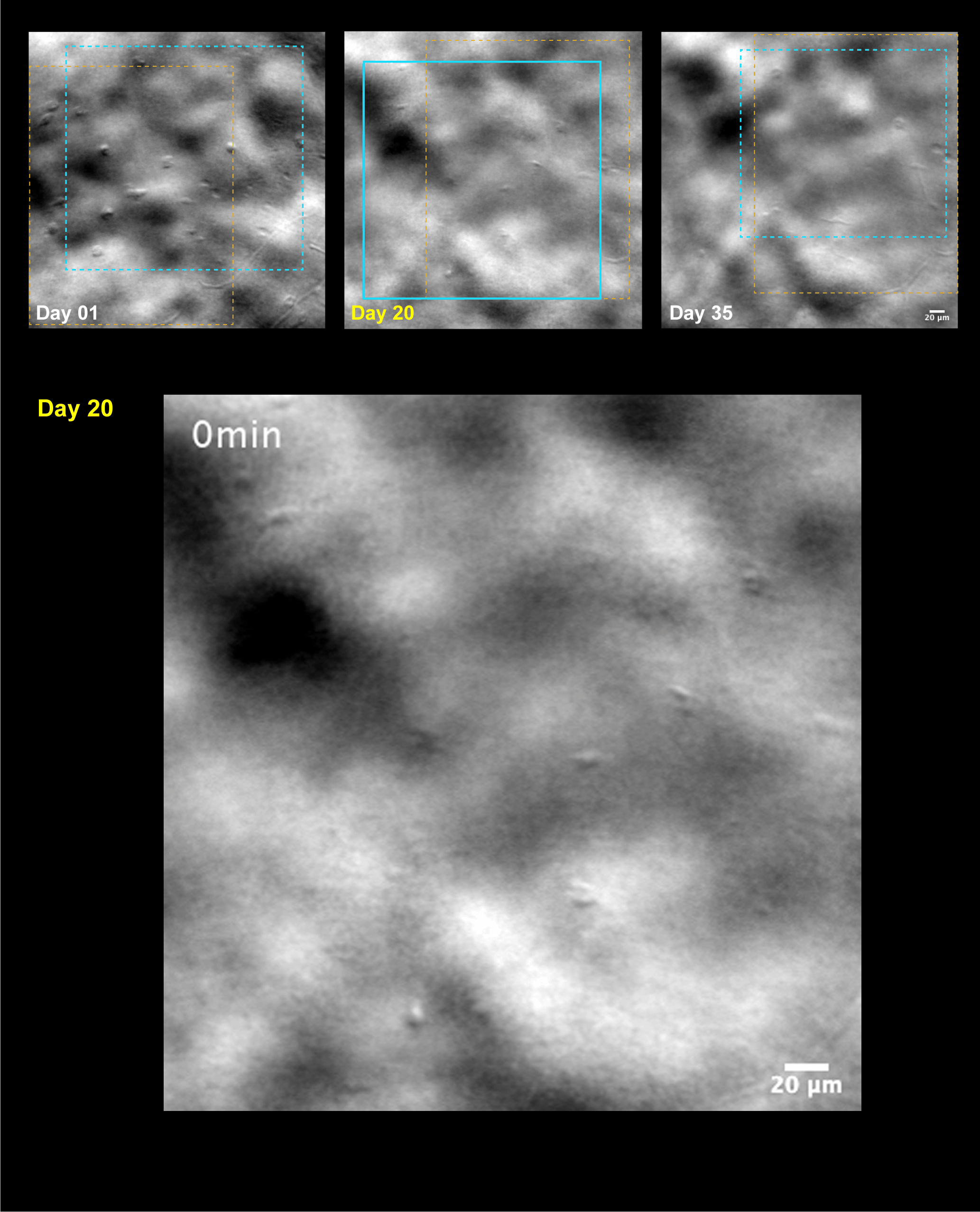

### immune cell motility observed in patient 1 with ASPPC on the third visit. Solid cyan boxes on top marked the region of time-lapsed below. Dashed cyan

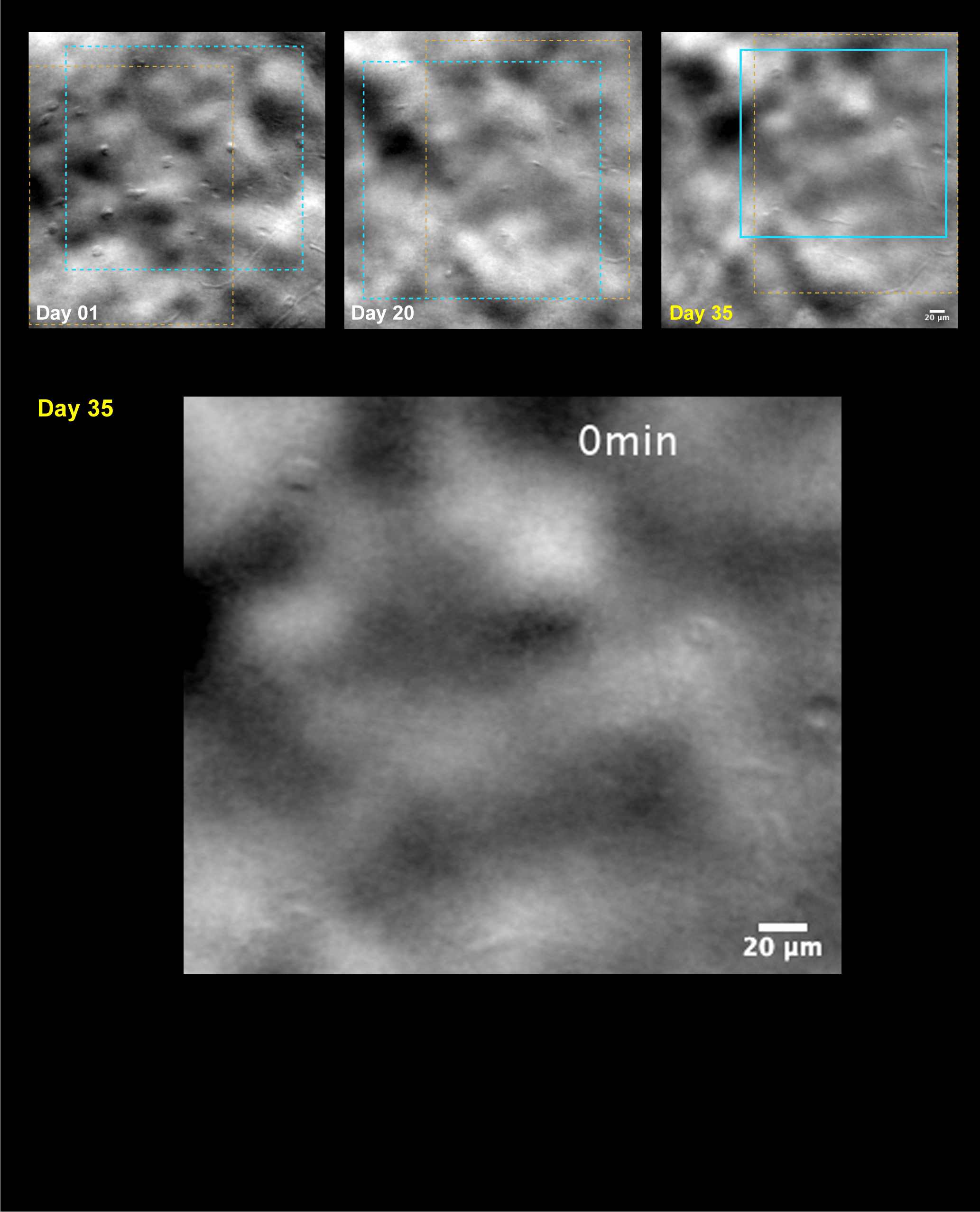

### Microglial cell motility observed in healthy retina (participant 1) with 30second intervals and corresponds to Fig.5A. White arrowhead tracks the posi

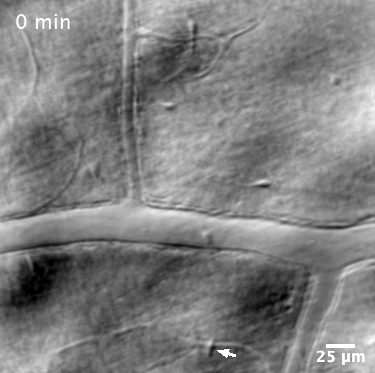

### Microglial cell motility observed in healthy retina (participant 1) with random intervals.

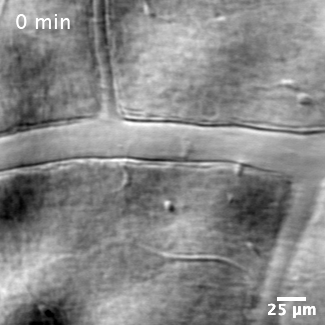

### Microglial cell motility observed in healthy retina (participant 2). White arrowhead tracks the cell that changes morphology locally.

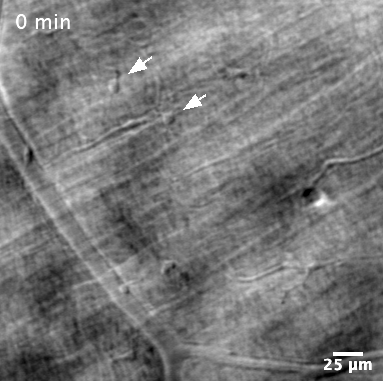

### Microglial cell motility observed in healthy retina (participant 3) and corresponds to Fig.5E. White arrowhead tracks the cell that changes morphology

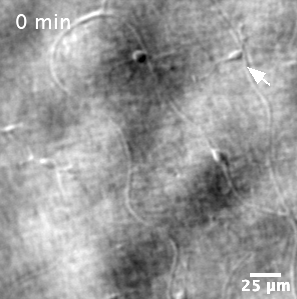

### Microglial cell motility observed in healthy retina (participant 5) and corresponds to Fig.5F.

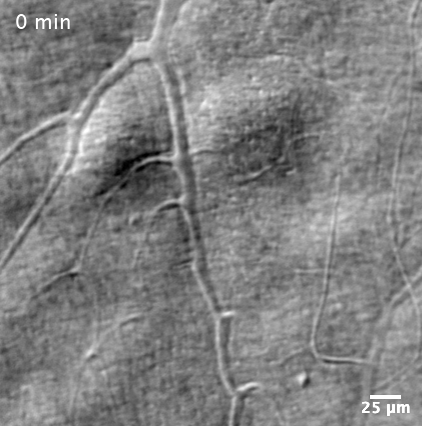

### Microglial cell motility observed in patient 2 with active uveitis. Microglial cells have relatively more activity by changing its morphology and posi

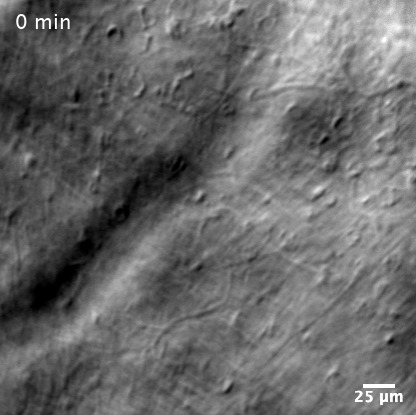

### Microglial cell motility observed in patient 3 with chronic uveitis. Microglial cells barely moved. Correspond to Fig.9C.

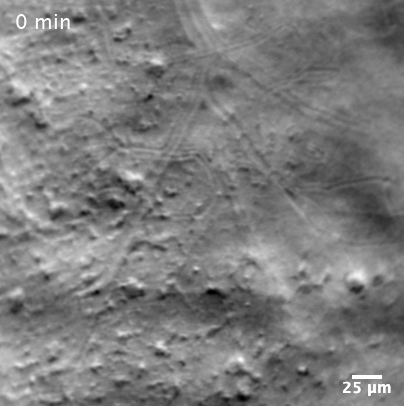
